# A generalised mechano-kinetic model for use in multiscale simulation protocols

**DOI:** 10.1101/2020.11.17.386524

**Authors:** Benjamin S. Hanson, Lorna Dougan, Oliver G. Harlen, Sarah A. Harris, Daniel J. Read

## Abstract

Many biophysical systems and proteins undergo mesoscopic conformational changes in order to perform their biological function. However, these conformational changes often result from a cascade of atomistic interactions within a much larger overall object. For simulations of such systems, the computational cost of retaining high-resolution structural and dynamical information whilst at the same time observing large scale motions over long times is high. Furthermore, this cost is only going to increase as ever larger biological objects are observed experimentally at high resolution.

We derive a generalised mechano-kinetic simulation framework which aims to compensate for these disparate time scales, capable of dynamically exploring a defined mechanical energy landscape whilst at the same time performing kinetic transitions between discretely defined states. With insight from the theory of Markov state models and transition state theory, the framework models continuous dynamical motion at coarse-grained length scales whilst simultaneously including fine-detail, chemical reactions or conformational changes implicitly using a kinetic model. The kinetic model is coupled to the dynamic model in a highly generalised manner, such that kinetic transition rates are continuously modulated by the dynamic simulation. Further, it can be applied to any defined continuous energy landscape, and hence, any existing dynamic simulation framework. We present a series of analytical examples to validate the framework, and showcase its capabilities for studying more complex systems by simulating protein unfolding via single-molecule force spectroscopy on an atomic force microscope.

**Author summary:** Our intention with this work is to provide a generalised, highly coarse-grained model to allow kinetic processes (conformational changes, protein unfolding, chemical reactions etc) to occur within the context of a dynamic simulation. Performing computationally intensive dynamic simulations to obtain kinetic information can be unjustifiably costly and scientifically inefficient, and so we instead want to emphasise that experimentally available kinetic information can be used to infer the underlying dynamics they result from. We hope our work can begin a discussion on the topic of computationally efficient science, and continue the drive towards collaborative science between theory and experimentation.

## Introduction

Over the past decade, experimental techniques have matured to such an extent that we can observe biological systems on length scales never before seen. Cryo-electron microscopy is enabling us to determine the atomistic structures of macromolecular complexes without the need for crystallisation, providing us with significantly more realistic instances of biological functional pathways [1–3]. Additionally, the continued development of various forms of super resolution microscopy and high throughput data analysis allow us to observe motions over ever shorter time scales [4, 5]. Many of these structures are being deposited into the Electron Microscopy Data Bank (EMDB), which has seen an almost exponential growth rate over the past decade [6]. Simulation techniques are being developed which utilise this data, especially at coarse-grained scales [7]. However, further consideration is needed with respect to the meaning of physical models at these newly explored spatio-temporal scales [8], and to the practicalities of performing computer simulations of these almost microscopic objects.

We may take as an illustrative example the molecular motor dynein, a system we have previously investigated [9, 10]. We consider that the features we outline now for dynein are common to many biomolecules, and to attempts to capture their operation in a multiscale simulation framework. Hanson *et al*. recently used existing all-atom molecular dynamics simulations of a single monomer of the motor to parameterise a much coarser continuum mechanical model of the same object [10], and found that a significantly coarser model can be used successfully for simulating the equilibrium dynamics. However, the biological function of the motor is in its so-called “powerstroke”, a change of equilibrium shape driven by cascading atomistic processes [11] which generates the directed force required for the motor to do the useful physical work [12–14]. Experiments show that the time between individual powerstroke events is typically much longer than either the mechanical relaxation time of the molecule or the time over which the powerstroke itself occurs. Hence the change in equilibrium configuration associated with the powerstroke is often considered as an effectively instantaneous chemical kinetic event [15]. However, given that the underlying structure of the motor [10] and its physical environment [9] affect its equilibrium, these kinetic events may not be completely independent of the underlying dynamics from which they emerge. The question then arises as to how one should include the powerstroke event into coarse-grained dynamical models of dynein, and reconcile these disparate timescales.

Dynein is not the only example of such disparate timescales in biological physics. We believe that the issues encountered above are sufficiently general to warrant consideration. Any coarse-grained biomolecular simulation might need to model transitions between different states, in which the configuration of a molecule at mechanical equilibrium is changed. Two particular examples of this are (i) molecular motors (as in dynein above) in which the configurational change between pre- and post-powerstroke states is driven by reactions within chemical-kinetic cycle, or (ii) protein unfolding, either at equilibrium [16–18] or driven by application of external force. External force can play a role in both of these examples. Translational molecular motors such as dynein have measurable stall forces [14, 19], and these forces can be applied by other molecular motors acting in the opposite direction [20]. Proteins are subjected to large forces in the artificial setting of single-molecule force spectroscopy experiments using an atomic force microscope (SMFS-AFM), but also in natural settings such as titin within muscle fibres [21], and engineered biological matrices such as protein hydrogels [22]. These external factors change the effective activation energy of the kinetic transition, and hence change the rate and manner by which thermal fluctuations and other dynamical effects drive transitions in these types of system.

The above discussion shows a need for a framework in which the disparity between dynamic and kinetic timescales are somehow reconciled, and their coupling explicitly modelled. This has been previously attempted specifically with respect to both molecular motors [23, 24] and much more commonly to protein unfolding [18, 25–30], but to our knowledge, a fully general approach has yet to be discussed. To address this considerable challenge in mesoscale modelling, we have developed a generalised ‘mechano-kinetic’ simulation framework which utilises both explicit mechanical modelling at mesoscopic length scales, whilst at the same time accounting for chemical-kinetic events beyond the scope of the dynamic model with a coupled kinetic model similar to a Markov state model. This coupling is such that the kinetic rates themselves dynamically alter as the system moves through the dynamic energy landscape and hence, using only the principle of detailed balance, our simulation framework enables the mesoscopic dynamical simulation to instantaneously contribute towards kinetic events such as conformational changes and chemical reactions, and thus implicitly affect dynamics at smaller length scales and vice versa.

## Theoretical Development

Our goal is to derive a ‘mechano-kinetic’ simulation framework; a simulation that is able to dynamically explore a defined mechanical energy landscape whilst at the same time perform kinetic transitions between discretely defined states. Here, and in what follows, we use the word *dynamics* to refer to changes in shape and conformation of a molecule as it explores a mechanical energy landscape; we use the word *kinetics* to refer to (discrete) changes of state which would alter that landscape (e.g. due to a chemical change such as a binding or unbinding event). We recognise it is usually not possible to cleanly distinguish these. Nevertheless, such a distinction is forced on us in coarse-grained modelling: “dynamical” changes refer to changes in molecular configuration explicitly resolved in the model, whilst “kinetic” changes are driven by processes at smaller, unresolved, scales.

### Mesostates

Importantly, we want the location in dynamical phase space to affect the kinetic processes in a thermodynamically consistent manner, such that we gain insight into how kinetic processes interact with different mechanical energy environments. To emphasise the target application of the biological mesoscale, and also to distinguish the discrete kinetic regime from continuous mechanical regime, we will refer to the discrete kinetic states as “mesostates”.

Fig.1 shows a schematic of a kinetic network, where each mesostate has some associated occupation probability *P_i_*. For a physical system, these occupation probabilities can be written in terms of a free energy, *F_i_*

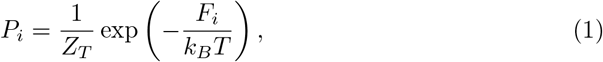

where *Z_T_* is the total partition function of the entire network, here acting as a normalisation constant. These probabilities, at equilibrium, must be related to the transition rates between mesostates, *R_ij_*, by the principle of detailed balance

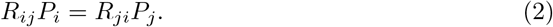

In principle, these probabilites and rates can be measured, either by experimentation or by a Markov State Model (MSM) calculation from a higher resolution simulation.

**Fig 1.**
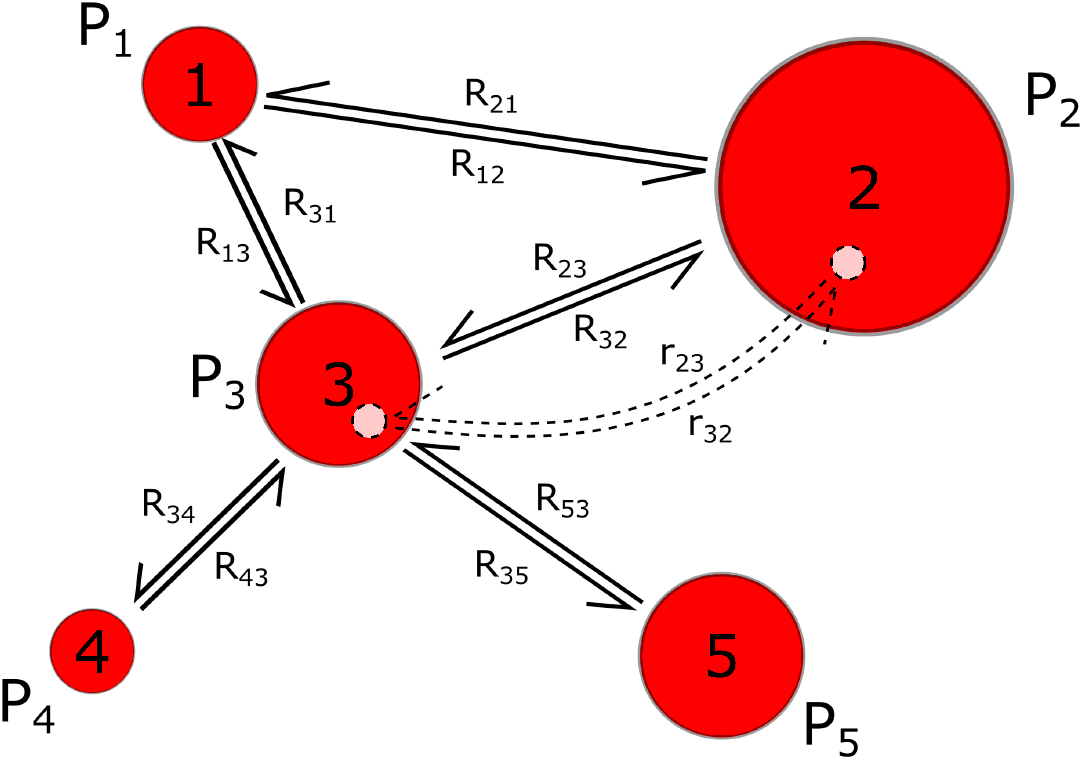
A dimensionless representation of a kinetic network with the number of states *N* = 5. Each state has an associated probability *P_i_* which is determined by the number of available microstates within it. Transitions between states occur explicitly between microstates with rates *r_ij_*, as shown by the dashed arrows, which then determine the relative transition rates *R_ij_* between each pair of states *i, j*, as given by the principle of detailed balance.

### Microstates and Free Energy

As shown for mesostates 2 and 3 in Fig.1, each mesostate contains within it a (potentially large number) of microstates. What these microstates are, and how they are parameterised, will depend on the system in question. In the examples considered here, the microstates are considered to be the set of (mechanical) shapes or configurations that a protein can take: we denote the microstate by the multidimensional vector 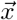. So, our physical picture is that the molecule exists within some mesostate *i* for some time, dynamically exploring changes of shape represented by changes in 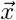.

The total partition function for the system is:

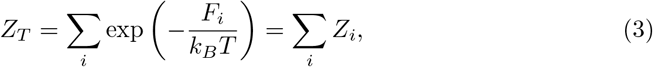

where the partition function for mesostate *i* is:

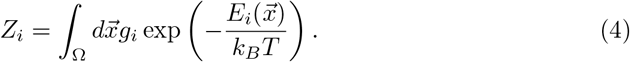

Here, 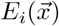 is the energy as a function of 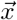 (it is, in fact, a free energy if 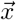 are coarse-grained variables). *g_i_* is the density of states within mesostate *i*. Any dependence of density of state on 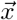 can be included as an entropic term within 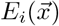 (and should be, since it will affect the dynamics of 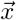). Then, the equilibrium probability density of system being at microstate 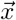 within mesostate *i* is:

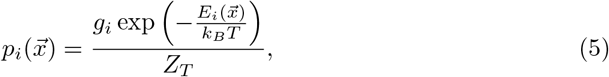

and 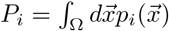. All dynamics and kinetics in the system must be consistent with these equilibrium distributions.

### Kinetic transitions between microstates

At some instant, a kinetic event may occur such that the molecule transitions from mesostate *i* to mesostate *j*. But, that transition must actually occur from a microstate within *i*, to a microstate within *j*, as illustrated explicitly for mesostates 2 and 3 in Fig.1. Detailed balance must also apply to such transitions between microstates.

We define 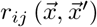 to be the transition rate from a single microstate at 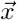 within *i* to a single microstate at 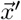 within *j*. Then, the equilibrium rate of transition from volume 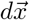 to volume 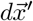 is the probability of being in 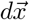, multiplied by the rate of transition per state, multiplied by the number of states in 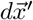:

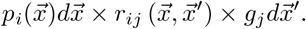

A similar equation applies to the equilibrium rate of reverse transition from 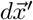 to volume 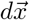, and hence (equating these two rates) we recover the expected detailed balance condition:

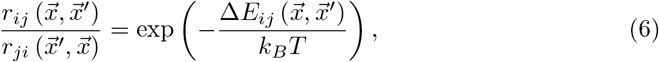

where 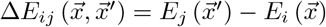.

Then, the total equilibrium rate of transition from mesostate *i* to mesostate *j* is obtained by integrating over all such pairs of volumes:

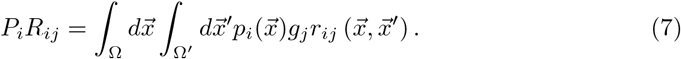

A similar equation applies to the reverse transition rate *P_j_R_ji_*. Applying Eq.6 thus ensures that Eq.2 is fulfilled. Eq.7 is useful to us in that it relates transition rates between microstates, 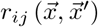, to the total transition rate between mesostates *R_ij_*.

A special case of the above is where the spaces 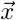 and 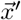 are identical, density of states *g_i_* = *g_j_* = *g*, and transitions occur only between the same point in phase space, i.e. 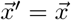 for all transitions. In this case we may write:

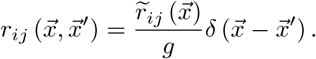

Then the microscopic detailed balance becomes:

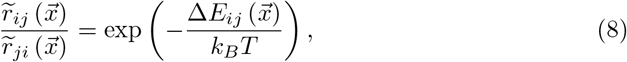

and Eq.7 simplifies as may be expected:

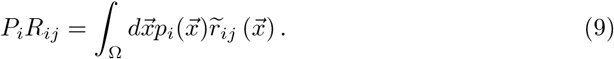

All the specific examples pursued in this paper are of this simpler form. The more general Eq.7 becomes necessary if transitions occur between different points of phase space within mesostates *i* and *j*, or if the phase spaces of *i* and *j* are not identical. The former may occur, for example, in models where a dynamic transition through phase space is explicitly defined as a part of a kinetic transition, and the latter in coarse grained modelling where the two mesostates are represented by different coarse grained models.

The above equations are extremely general. They relate forward and reverse transition rates (via detailed balance), and mesoscopic transition rates to microsopic transition rates. These equations place constraints on what is “allowed” in a model. However, they are not sufficient to fully constrain a model: one needs to make further physical assumptions. In what follows, we explore some possible assumptions that are consistent with the preceding work.

### Candidate microscopic models

We find it convenient to separate the energy of a microstate as 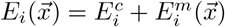, where 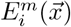 is the “mechanical” energy used to compute dynamics of 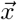 within mesostate *i*, and 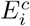 represents all other contributions to the (free) energy of mesostate *i* that are independent of 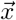. Correspondingly, energy differences can be written as 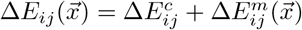.

This then allows us to write Eq.8 in the form

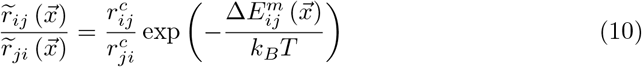

where the ‘intrinsic’ rates 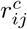 are related as

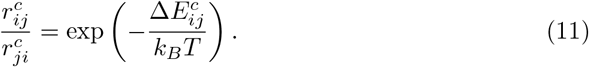

We note that these intrinsic rates are not necessarily independent of 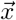: it may be, for example, that transitions are only possible within some region of phase space. However, their ratio is independent of 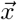. For the purposes of this work we will assume them to be constant, but recognise that so long as Eq.11 is satisfied, they can take any functional form.

We now indicate two candidate forms of model that are consistent with Eq.10.

1. **Mechanical energy difference model.** With insight from the work of Sarlah and Vilfan [23, 24], we separate the detailed balance condition of Eq.10 into individual rates as

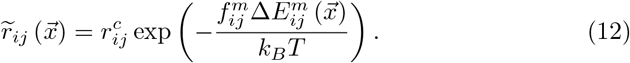 The factor 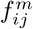 indicates the fraction of the mechanical energy difference 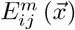 that influences the forward transition rate 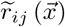, the remainder being attributed to the reverse transition. Eq.10 then requires the constraint 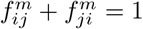. The implicit assumption is of an energy barrier to the transition, located at some position along the reaction co-ordinate between mesostates *i* and *j*. The mechanical energy alters the height of that barrier in an Arrhenius-like fashion. 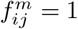 suggests the barrier is located close to state *j* along the reaction co-ordinate, whilst 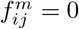 indicates the barrier is close to *i*. We can now relate Eq.12 to equilibrium transition rates at the mesoscale. Substitution into Eq.9, and using Eqs.1 and 5 with *g_i_* = *g* gives

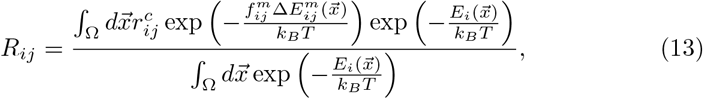

and, with the assumption of constant 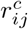, we finally obtain

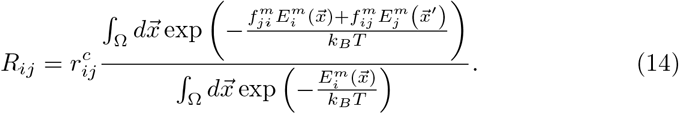 This equation determines the value of 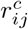 required in the model to obtain a given value of *R_ij_* when the system is at thermal equilibrium. So, for example, *R_ij_* may be an experimentally observed transition rate between two mesostates. Having now determined the appropriate choice of 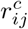, we can continue to use Eq.12, for example in a non-equilibrium situation where an additional external force is applied to the system, which would then result in a change in the overall average transition rate.
2. **Force-activated transition model.** A reasonable assumption in some situations is to consider that the transition rate from *i* to *j* is driven by some component of a force 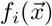, which depends upon the configuration 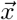. So, for example, 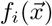 could be a force acting across a bond, which may dissociate to drive a change of state (e.g. a protein unfolding). This force might (in many cases) be obtained from the gradient of the mechanical energy 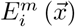 with respect to one of the components of 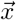. This distinguishes the force-activated transition from the mechanical energy difference model (where the transition rate is related only to the difference in energy between mesostate *i* and *j* rather than gradient of the mechanical energy within a mesostate). Models explicitly considering external force have long been used ubiquitously, most relevant to us in the case of protein unfolding [31, 32]. However, these models are often used in isolation at the free-energy (mesostate) level, whereas we show that external forces can be explicitly included at the mechanical (microstate) level in the same manner. If the critical extension of the bond for dissociation is a small distance *b*, where *b* = *b_ij_* = *b_ji_*, the energy barrier towards the transition is then reduced by 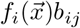 giving transition rate:

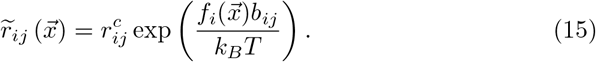 It may be that the rate of the reverse transition is negligible and can be ignored. However, if re-association is to be considered a possibility then Eq.10 requires that:

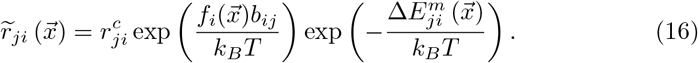 We can again relate these microscopic transition rates to equilibrium transition rates at the mesoscale. Substitution into Eq.9, and using Eqs.1 and 5 with *g_i_* = *g* gives, for the forward rates:

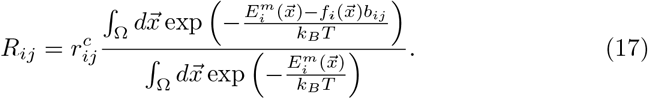 However, for the reverse rates:

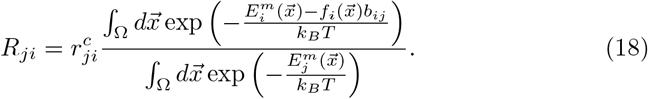 In an unbinding transition, for which this model is most appropriate, it would typically be the case that mesostate *i* (representing bound state) would be mechanically stiffer than mesostate *j*. Indeed, many proteins are held in their folded conformation by strong disulphide bonds in a variety of configurations [33], and it may be the extension of these bond complexes that lead to the value *b_ij_*. Hence, integrals over 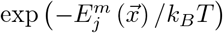 are dominated by relatively small region of phase space V_i_, whilst integrals over 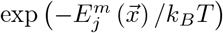 are dominated by a larger region of phase space *V_j_* ≫ *V_i_*. Thus, the reverse transition rate in Eq. 18 contains an implicit factor of *V_i_*/*V_j_*, representing the fact that rebinding can only occur from state *j* to state *i* if the system is found in the correct region of phase space, close to the favourable region for state *i*. If *V_j_* ≫ *V_i_* then such occasions will be rare, through random thermal fluctuation. We note here that the factor 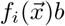 occurring in these equations is a linearised energy contribution, which will typically only be appropriate in the region *V_i_* of phase space where mesostate *i* undergoes typical fluctuations. It may be that it is appropriate to set 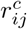 and 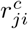 to zero outside this region (and taking it back inside the integrals), so as to prevent the factor 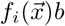 in equations 17 and 18 from creating spurious, unphysical effects.

The following section contains various examples which show the capabilities of the method, and hint at its generality and wide field of potential applicability.

## Results

Fig.2 shows the various systems investigated in this section. Before discussion of those, however, we begin by showing how Eq.14 can be calculated for a specific system, and how Eq.12 is applied as part of a dynamic simulation protocol. This is done using the mechanical energy difference model, although an equivalent set of calculations could be performed for the force-activated transition model using Eqs.15–18.

**Fig 2.**
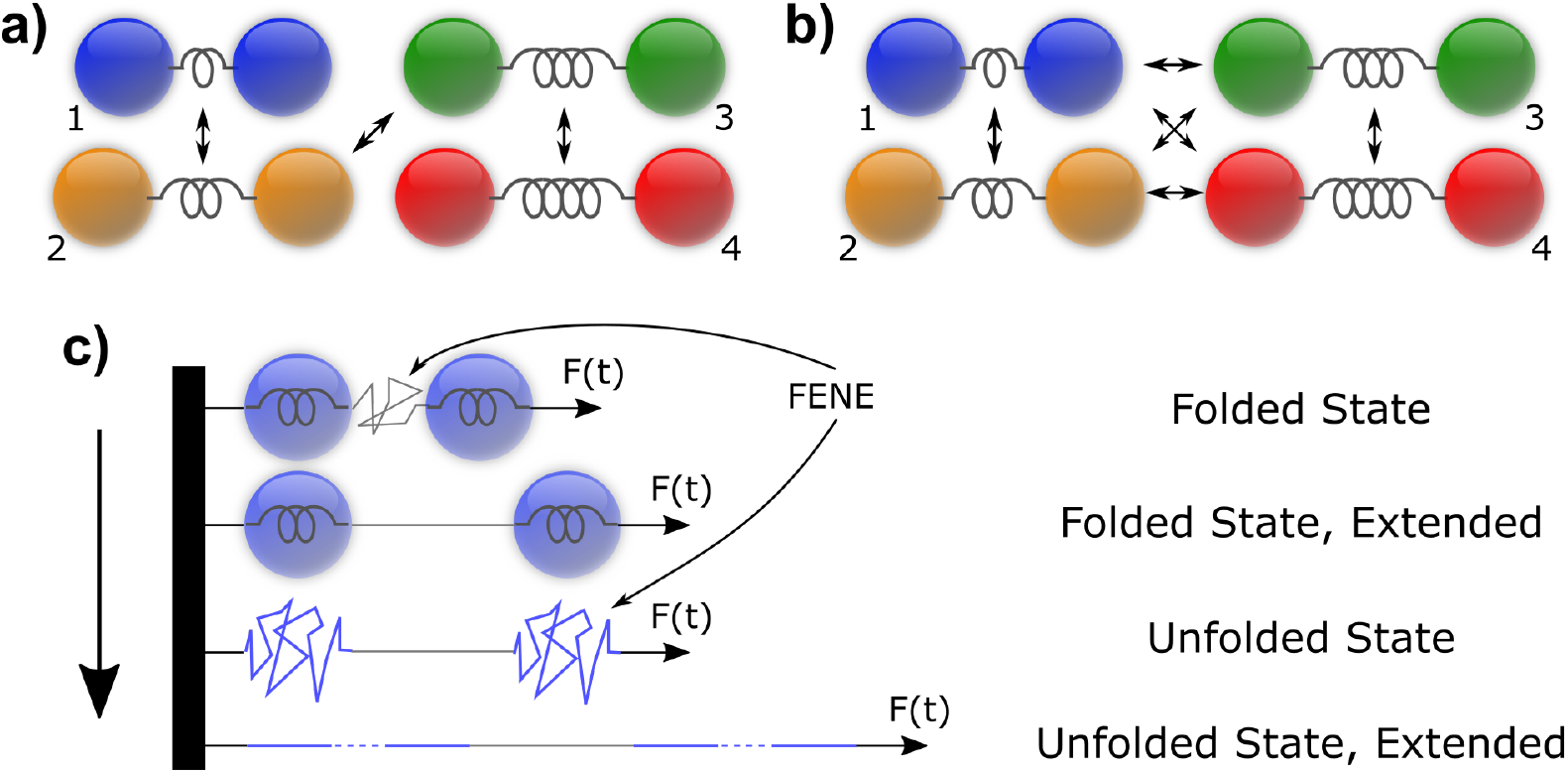
The structural models in each of our example systems. **a)** Four mesostates, defined by their different equilibrium lengths and spring constants and coloured to correspond to the associated mesostate graphs throughout this article. Here there is only a single transition pathway from mesostate 1 to mesostate 4, representing a process akin to (but distinct from) a reaction co-ordinate. **b)** Similar systems to **a)**, but this time every possible kinetic transition is allowed. **c)** By representing “proteins” as Hookean springs which can transition to springs with a modified finitely extensible non-linear elastic (FENE) potential, i.e. a polypeptide chain [32], we obtain a representation of protein unfolding by mimicking SMFS-AFM experiments.

## Applying the mechano-kinetic framework

To investigate the capabilities of the model, we begin with a simple 1D dumbbell model; two beads connected by a Hookean spring with spring constant *k_i_* and equilibrium length *l_i_*. The dynamic simulation framework we use is a Brownian dynamics protocol, numerical integration of the Langevin equation where inertia, and hence velocity degrees of freedom, are neglected. For the two dumbbell coordinates *x*_1_ and *x*_2_, the Langevin equations are

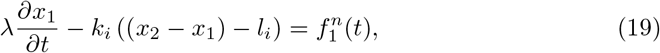

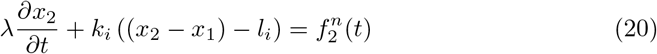

Each bead is subject to a Stokes drag *λ* and an associated stochastic thermal noise 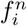 calculated via the fluctuation-dissipation theorem, with associated statistics

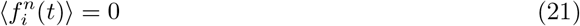

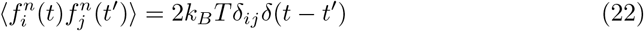

The corresponding mechanical energy for the system is given by

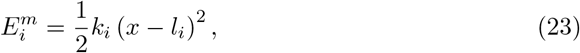

where *x* = *x*_2_ − *x*_1_ defines the single elastic degree of freedom within the system.

The kinetic simulation component utilises the sets of internal spring parameters, *k_i_* and *l_i_*. These two parameters define the mesostate, i.e. *k_i_* and *l_i_* are the parameters for mesostate *i*, and will vary between mesostates. When altered, these parameters change the energy landscape independently of the location of the dumbbell nodes within phase space, setting *x* = *x′* in Eq.14. In principle all parameters (e.g. the drag) could vary between mesostates, but for clarity we consider only kinetic alterations to the elastic energy landscape. In the simulations reported below, we set *λ* = 6*π*pN.ns/nm and remains constant across all mesostates.

With these independent definitions of the dynamic and kinetic simulation components, we can now couple them together. For this 1D system and in the absence of intertia, the mechanical energy term within Eq.14 is greatly simplified. Introducing Eq.23 into Eq.14 and expressing the denominator integral exponent as a polynomial in *x* gives only Gaussian integrals in *x* and so can be solved exactly, yielding

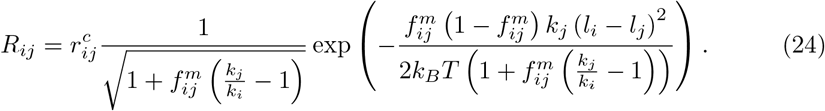

Eq.24 completes the coupling between dynamic and kinetic simulation components within the mechano-kinetic framework. From a set of *R_ij_* together with a dynamic model, we can calculate the set 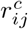. Now, following each dynamic simulation timestep, we include the probability of a mesostate transition as per Eq.12, where we convert the rates to transition probabilities by multiplying by the simulation timestep. Eq.12 therefore makes these transition probabilities explicitly dependent on the current microstate mechanical energy. If a transition occurs, we keep the two node positions constant, change the spring parameters, and allow the dynamic simulation to continue. If at any point a probability exceeds 1, we stop the simulation and lower the timestep to allow the correct resolution of kinetic events.

The introduction of instantaneous kinetic transitions within a dynamic simulation introduces the potential for non-convergence of statistical properties. Physical validation of each simulation component is provided as Supporting Information.

### A four-state system

To investigate the behaviour of the model when applied to a molecule undergoing conformational changes, we have designed a series of kinetic transitions using the dumbbell model corresponding to a four state system, in which the average length of the dumbbell increases throught the states. We define two alternative systems of this set of kinetic mesostates, given by their mechanical parameterisations in Table 1, together with an associated equilibrium kinetic transition rate matrix corresponding to both systems given in Table 2. The schematic of this system is shown in Fig.2a.

**Table 1.** The two families of material parameters used to define the mesostates in the four-state kinetic networks of the dumbbell model. The first (left) set contains equal and relatively low values of *k_i_* whereas the second (right) set of *k_i_* are much higher. The values of *l_i_* linearly increase with the mesostate index.

| Mesostate ($i$) | $k_i$ (pN/nm) | $l_i$(nm) | Mesostate ($i$) | $k_i$ (pN/nm) | $l_i$(nm) |
| --- | --- | --- | --- | --- | --- |
| 1 | 10 | 1 | 1 | 200 | 1 |
| 2 | 10 | 2 | 2 | 50 | 2 |
| 3 | 10 | 3 | 3 | 50 | 3 |
| 4 | 10 | 4 | 4 | 200 | 4 |

**Table 2.** The average transition rates between mesostates used in the four-state kinetic networks of the dumbbell model. The mesostate occupation probabilities have been analytically calculated for comparison with simulation.

| Transition Rates, $R_{ij}$ ($\mu\text{s}^{-1}$) | | | | | $P_i$ |
| --- | --- | --- | --- | --- | --- |
| From / To | 1 | 2 | 3 | 4 |  |
| 1 | N/A | 0.916 | - | - | 0.013 |
| 2 | 0.337 | N/A | 50 | - | 0.035 |
| 3 | - | 6.767 | N/A | 18.394 | 0.256 |
| 4 | - | - | 6.767 | N/A | 0.696 |

Since Table 2 defines all of the expected transition rates, which in principle may have been experimentally measured, it also implicitly defines the mesoscale free energy landscape and all of the resultant occupation probabilities. Thus our investigation will tell us nothing new at the mesoscale level at this stage. Instead, our aim is to analyse the modifications to the kinetic rates made by the continuous underlying dynamics, and verify that the measured, average kinetic behaviour is reproduced. Note that many of the transition rates shown in Table 2 are zero, indicating that direct transitions cannot occur between these states. However, transitions via other intermediate states can still occur (i.e. from mesostate 1 to mesostate 3 via mesostate 2).

The energy landscapes implied by this kinetic system are illustrated in Fig.3. We see in Fig.3a) that as all of the stiffness parameters *k_i_* are equal in the first parameterisation, no particular state is preferable over long periods of time with respect to mechanical energy. On the other hand, in the second parameterisation we include two high stiffness states (*k*_1_ and *k*_4_) and two low stiffness states (*k*_2_ and *k*_3_). However, irrespective of the mechanical energy environment, the transition rates in Table 2 imply certain characteristics of the free energy landscape along a “pseudo-reaction co-ordinate” between mesostates. If *R_ij_* > *R_ji_*, then this implies (via Eqs.1 and 2) that the free energy of state *i* is higher than state *j*. Smaller overall values of *R_ij_* and *R_ji_* typically imply larger energy barriers between the two states. We have set 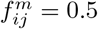 for all *i*, *j* in the simulations, which implies the energy barrier lies half-way between the two states along the reaction co-ordinate. Given this, the transition rates in Table 2 imply a one-dimensional reaction co-ordinate of the form shown in Fig.3. We emphasise that this “pseudo-reaction co-ordinate” is not the extension *x* of the dumbbell, but rather some underlying reaction variable along which lie our implicitly coarse-grained kinetic states.

**Fig 3.**
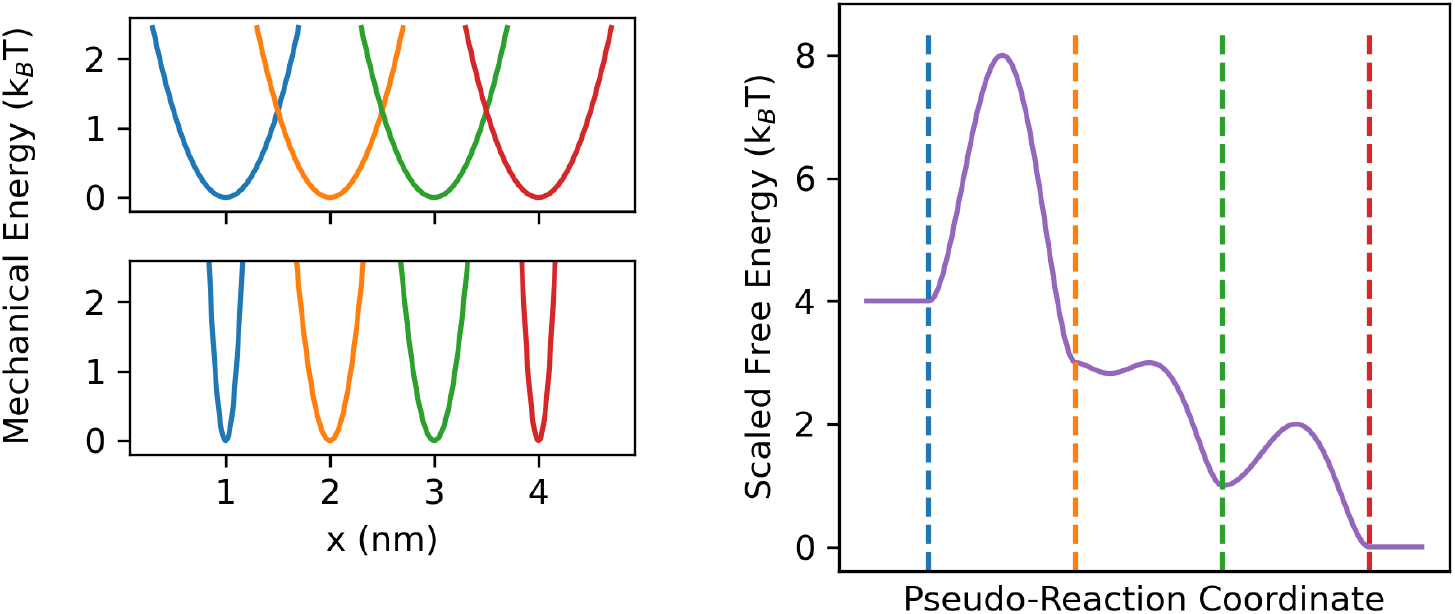
The two “levels” of energy landscape within the four-state kinetic networks of the dumbbell model. a) The mechanical energy landscapes given in Table 1, where the occupation of each state is given a different colour. Top: Set 1 - All values of *k_i_* are equal. Bottom: Set 2 - Values of *k_i_* vary between mesostates. b) A possible representation of the free energy as a function of some hypothetical “pseudo-reaction co-ordinate”, with barriers implied by the rates given in Table 2. The “center” of each mesostate is shown as a vertical dashed line.

#### Four state system - Results

For each mechanical simulation parameterisation, Table 3 shows the rates 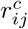 which are modified by exploration of the mechanical energy landscape to obtain the *R_ij_* values shown in Table 2. Let us briefly investigate some of the properties of these values. For the set of 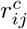 where all the *k_i_* are equal for the four mesostates, each 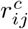 is increased from its corresponding *R_ij_* value by the same multiplicative factor. This is because the mechanical energy environments for each state are identical, albeit translated in space, and so with the application of Eq.24, all integral results are the same for every state pair. The presence of a mechanical energy landscape here, then, acts as a universal barrier to kinetic transition, such that an extra amount of energy is required to make the kinetic transition. For the other set, corresponding to the mesostates with differing *k_i_* values, we see a similar effect for 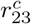 and 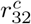 which, having the same mechanical energy landscape, are increased from their corresponding mesoscopic values by the same factor. The increase here is greater than in the previous (equal *k_i_*) network, as the values of *k_i_* are significantly higher. Importantly, because the values are increased by the same factor, the detailed balance condition between the pair of states is unchanged by the presence of a mechanical energy landscape and thus, while that rates are slower, the contributions to the equilibrium probabilities will be the same. However, this is not the case for any of the other pairs of rates in this network. As the mechanical energy environment changes between mesostates 1 & 2, and 3 & 4, the relative increase of 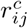 from the associated *R_ij_* value is different, showing that the detailed balance condition of the 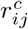 values are not the same as the mesoscopic *R_ij_* values. In other words, dynamic motion through a mechanical energy landscape, in the general case, modifies the detailed balance conditions between the pre-defined mesostates in our model. In reality the reverse is true, in that the presence of a mechanical energy landscape alters the implicit detailed balance conditions between microstates, leading to a different detailed balance condition at the free energy level.

**Table 3.** The instantaneous transition rates between mesostates used in the four-state kinetic networks of the dumbbell model corresponding to the two sets of mesostates given in Table 1. Top: Equal, low *k_i_* mesostates. Bottom: Differing, large *k_i_* values.

| Transition Rates, $r_{ij}^c$ ($\mu s^{-1}$) | | | | |
| --- | --- | --- | --- | --- |
| From / To | 1 | 2 | 3 | 4 |
| 1 | N/A | 1.241 | - | - |
| 2 | 0.457 | N/A | 67.761 | - |
| 3 | - | 9.171 | N/A | 24.928 |
| 4 | - | - | 9.171 | N/A |
| From / To | 1 | 2 | 3 | 4 |
| 1 | N/A | 8.239 | - | - |
| 2 | 6.063 | N/A | 228.56 | - |
| 3 | - | 30.934 | N/A | 330.91 |
| 4 | - | - | 60.869 | N/A |

Although the 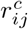 values calculated and used in each simulation are different, Table 4 shows the equilibrium mesostate occupation probabilities for both sets of mechanical material parameterisations, calculated by averaging the expected occupation probabilities from a single, long-time simulation over many repeats beginning in different initial states. We can see that the mesostate occupation probabilities for the two simulations are equivalent to the analytically calculated values given in Table 2, despite the large differences in the values of *k_i_*. This clearly shows that the mesostate occupation probabilities within our model do indeed remain constant as required, being fixed by the pre-defined *R_ij_* values, whilst the underlying mechanical energy landscape is modified.

**Table 4.** The final expected mesostate occupation probabilities within the four-state kinetic networks of the dumbbell model, which accurately reproduce the analytically calculated rates given in Table 2. Each set corresponds to a different set of material parameters as defined in Table 1. Uncertainties in the last digit are shown in brackets.

| Mesostate | Mesostate Occupation Probabilities, $P_i$ | | | |
| --- | --- | --- | --- | --- |
|  | 1 | 2 | 3 | 4 |
| Set 1 | 0.012(1) | 0.034(1) | 0.256(2) | 0.698(2) |
| Set 2 | 0.014(1) | 0.036(1) | 0.258(2) | 0.693(3) |

Fig.4 shows the probability density of length fluctuations at equilibrium from a single simulation of each set of mesostates. We see that whilst the kinetic rate modifications within the model take the mechanical energies into account in order to return the average rates *R_ij_*, the mechanical energy landscape is still fully realised within the framework. In particular, we see that the low mechanical stiffness of individual states in Fig.4a) means that length fluctuations within each state are relatively large compared to the associated equilibrium lengths, resulting in a unimodal (albeit skewed) overall distribution centred around, but not exactly on, the most likely mesostate. Within a standard Markov state model, each of these states would likely be indistinguishable from one another, yet our implied knowledge of both the mesostate kinetics and mechanical energy landscape allows us to include them anyway, such that their indistinguishability is effectively a deduced property. On the other hand, for the second parameterisation in Fig.4b), the higher mechanical stiffness suppresses the length fluctuations within each mesostate so that the individual mesostates are distinguishable in the overall length distributions, with clear Gaussian distributions centered on each equilibrium length. This is less apparent for mesostates 1 and 2, however, as their associated free energies are significantly higher, and hence their occupation probabilities lower.

**Fig 4.**
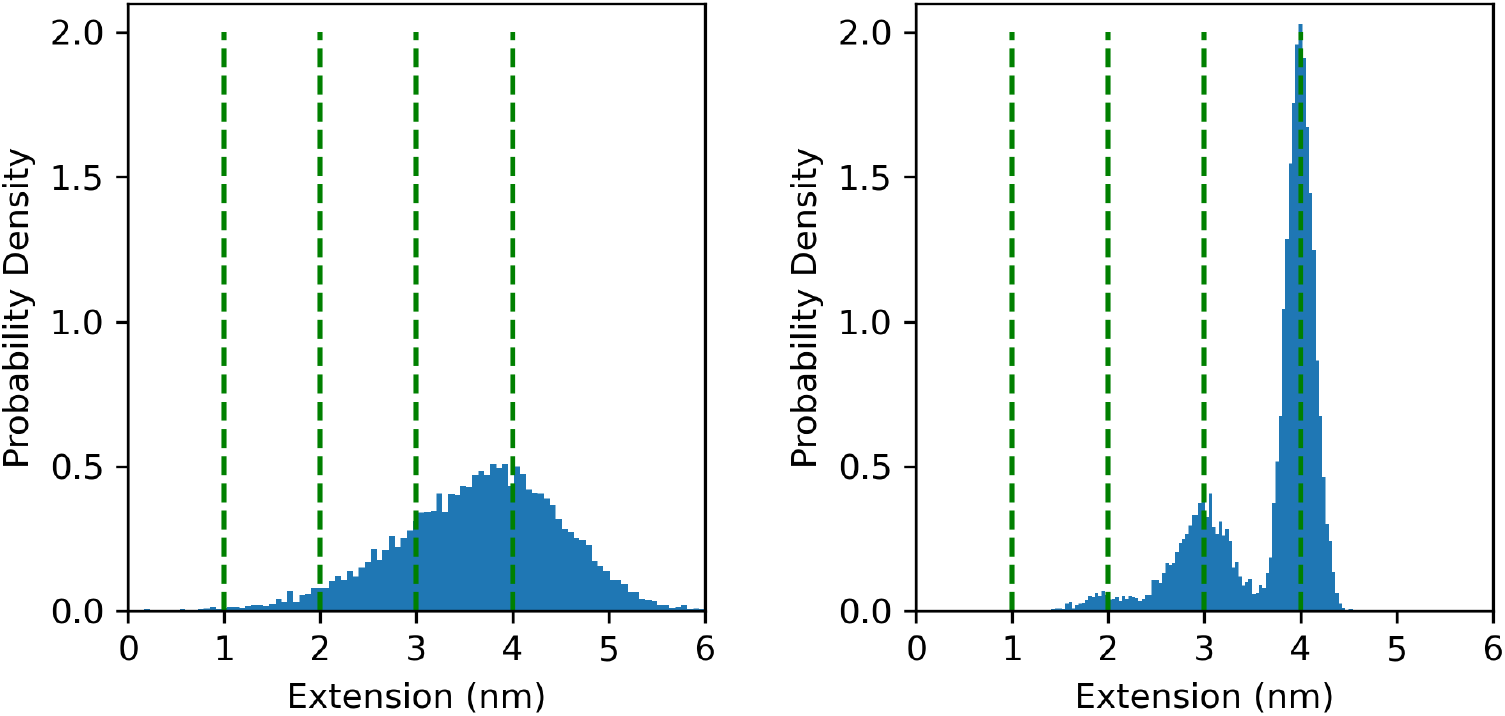
The microstate probability densities from simulation of the four-state kinetic networks of the dumbbell model. The mechanical equilibrium positions corresponding to each state are shown as dashed green lines. a) Mesostate set 1 b) Mesostate set 2.

Overall we see that within the mechano-kinetic framework, no matter the structure of the mechanical energy landscape, the introduction of Eq.24 into Eq.12 enables the simulation to reproduce the *R_ij_* values used as input parameters. As such, we have used the measured kinetic rates given in Table 2 together with the spring parameters in Table 1 to infer the specific form of the mechanical energy landscape. However, we will see in the next section that we can go further than the simple reproduction of existing measurements, and consider how direct external modifications to the mechanical energy landscape affect the resultant kinetic behaviour.

### External Forces

We now investigate how this framework behaves when an external potential is added to the equilibrium system. As previously discussed, we may wish to determine the effect of applying external mechanical forces to biological objects such as proteins or molecular motors. Consider the case where either experiments or high-resolution simulations have been performed and both the free energy and mechanical energy landscapes of the object in isolation are well known. For the case of dynein, the illustrative motor protein discussed in the introduction, both mechanical [34, 35] and kinetic [12, 36] information is available to parameterise the system. For proteins and unfolding in general, Hughes *et al*. provide a detailed review [32]. Using this information, we can use the mechano-kinetic simulation framework to make predictions based on these experimental systems.

To explore this type of problem in a general sense, we have designed two kinetic networks, both of which use the dumbbell model together with set of mesostates defined in Table 5. The first network is similar to the previous four-state example, whereas the second network is a fully connected kinetic network, where direct transitions between all states are possible. The schematic of this second system is shown in Fig.2d. The associated transition rates defining these networks are given in Table 6, top and bottom tables respectively. In contrast to the previous unforced four-state example, we here observe that because all of the rates are the same, when unperturbed, this system initially ought to have an equal probability of being in each of the mesostates. However, under the subsequent application of external forces we would expect the forces, as a source of mechanical energy, to bias the states in some manner. Within the *dynamic* simulation component, we apply a constant external force *f^ex^* to both nodes of the dumbbell but in opposite directions, therefore biasing the dumbbell towards higher extensions. To clarify, we use the values of 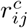 as given at equilibrium in the absence of external forces, and modify the reaction rates via Eq.12 to account for additional contributions to the mechanical energy from the external forces. With this approach we are able to use the mechano-kinetic model to predict not only how the application of this external force within the dynamical simulation affects the overall kinetic transition rates between mesostates, but also determine where those transitions are likely to occur in the mechanical energy landscape.

**Table 5.** The material parameters used to define the mesostates used in the kinetic network of the dumbbell model with external forces applied.

| Mesostate ($i$) | $k_i$ (pN/nm) | $l_i$(nm) |
| --- | --- | --- |
| 1 | 25 | 1 |
| 2 | 25 | 2 |
| 3 | 25 | 3 |
| 4 | 25 | 4 |

**Table 6.** The matrices defining the two sets of average transition rates between mesostates, together with the expected mesostate occupation probabilities, used in the kinetic network of the dumbbell model with external forces applied. Top: A single path through the mesostate network. Bottom: A fully connected kinetic network. These rates were chosen such that the total rate of leaving any given mesostate is equivalent to that in the single pathway model.

| Transition Rates, $R_{ij}$ ($\mu s^{-1}$) | | | | | $P_i$ |
| --- | --- | --- | --- | --- | --- |
| From / To | 1 | 2 | 3 | 4 |  |
| 1 | N/A | 50 | - | - | 0.25 |
| 2 | 50 | N/A | 50 | - | 0.25 |
| 3 | - | 50 | N/A | 50 | 0.25 |
| 4 | - | - | 50 | N/A | 0.25 |
| From / To | 1 | 2 | 3 | 4 |  |
| 1 | N/A | 16.6 | 16.6 | 16.6 | 0.25 |
| 2 | 16.6 | N/A | 16.6 | 16.6 | 0.25 |
| 3 | 16.6 | 16.6 | N/A | 16.6 | 0.25 |
| 4 | 16.6 | 16.6 | 16.6 | N/A | 0.25 |

#### External Forces - Results

Fig.5 shows the steady state occupation probabilities of each mesostate as a function of applied force, for both sets of rates. These were calculated by averaging the expected occupation probabilities from a single, long-time simulation over three repeats, with each simulation beginning in mesostate 1 at mechanical equilibrium. Although all of the values of *k_i_* in Table 5 and initial transition rates in Table 6 are equal, indicating no native energetic preference for any of the mesostates (as can be seen when zero external force is applied), the application of force immediately biases the system towards larger extensions. We also see from Fig.5b) that the single pathway and fully connected networks are almost identical at the kinetic scale. For each mesostate in ascending order, the root-mean-square deviations of the residuals of one graph with respect to the other are 0.006, 0.004, 0.006 and 0.006 respectively. This indicates that the steady-state occupation probabilities as a function of applied force are independent of the connectivity of the kinetic network, given simply by the Boltzmann distribution modified to account for the extra energy contribution from the externally applied force. Detailed balance implies that this change is brought about within the model via a change in the resulting values of *R_ij_*, but key is that this is propagated from the dynamic component of the model into the kinetic component.

**Fig 5.**
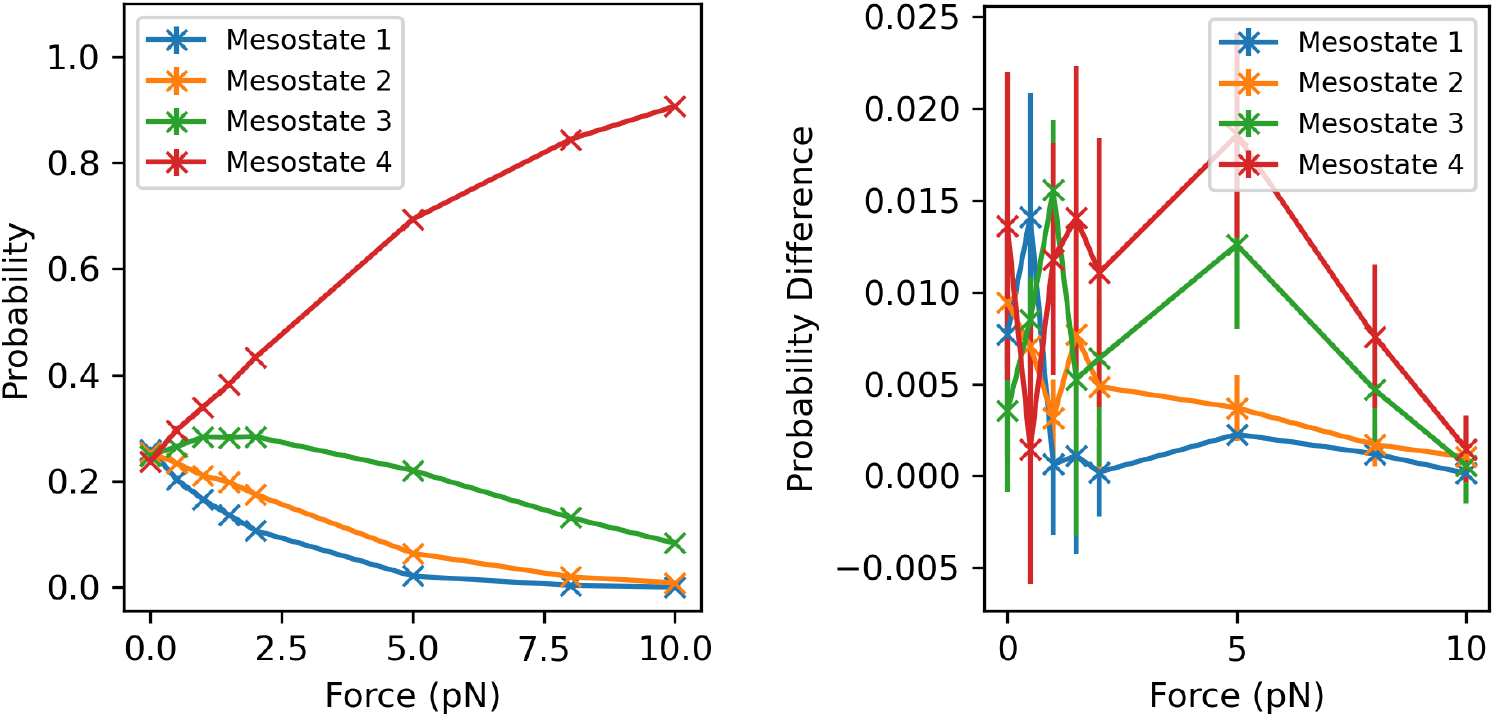
The expected occupation probability of each mesostate of the dumbbell model as a function of applied force. a) The single pathway network. b) The difference between the single pathway and fully connected networks. Note that, as expected, these traces are therefore identical for the two networks.

The application of force across all mesostates effectively generates an additional contribution to the free energy. As such, these forces can be included in pure kinetic models as free energies, 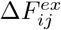, using a term such as 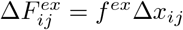 where Δ*x_ij_* is some approximation of the expected extension. This is similar to the force-activated transition model, except that the force is introduced at the kinetic (free-energy) level. However, with the explicit inclusion of the mechanical energy environment, we can observe the subtleties occurring at the microscale. Fig.6 shows the probability distributions, in the single pathway system, of dumbbell extension for zero external force and 10pN respectively, separated by mesostate. Overlapping these in a lighter colour are the distributions of dumbbell extension at the moment of transition into another mesostate. We see Gaussian distributions in the extensions, indicating thermodynamic equilibrium and reinforcing the results of Fig.5. In the Fig.6a) transition distributions, we see either single or bimodal overlapping Gaussian distributions centred on the the half-way points between the each neighbouring pair of states, a direct consequence of the fact that the difference between two quadratic energy potentials is itself quadratic. Under external force, however, in Fig.6b), we observe all of these probability distributions have effectively been pulled over to higher extensions, with the bimodal distributions of mesostates 2 and 3 forming a single mode centred at higher extensions, and mesostate 1 having a much smaller occupation and number of transitions occurring.

**Fig 6.**
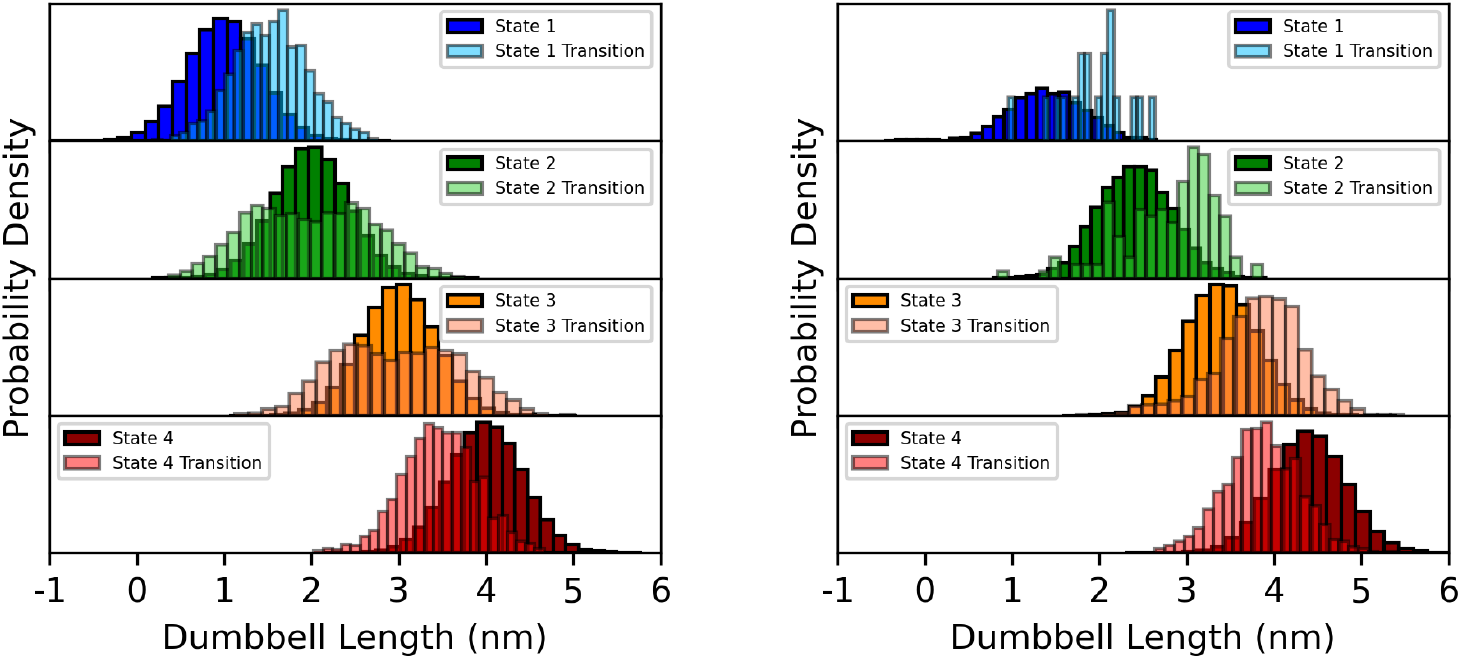
The normalised probability distributions, in the single network pathway system, of the dumbbell node separations and the separations specifically at the moment of transition into another mesostate. a) 0pN force applied. b) 10pN force applied.

Fig.7 shows the same information but for the fully connected network system. We observe approximately the same occupation probability distributions as in Fig.6, as may be expected from Fig.5, but the transition probability distributions are slightly more spread out. This is especially apparent if we compare mesostate 1 of Fig.6a) and Fig.7a), and mesostate 4, and is due to the fact that the fully connected network has more possible transition pathways. Indeed, as defined by the rates (in the absence of external force), mesostate 1 has an equal probability of transitioning straight to state 4 on average as it does mesostate 2. We therefore see that while the two systems shown yield the same occupation probabilities per mesostate, and change similarly under the application of force external to the initial parameterisation using Eq.24, the dynamical motion through phase space that we infer from the application of force at the continuous microstate level is dependent upon the structure and connectivity of the the kinetic landscape, i.e. which kinetic transitions are permitted.

**Fig 7.**
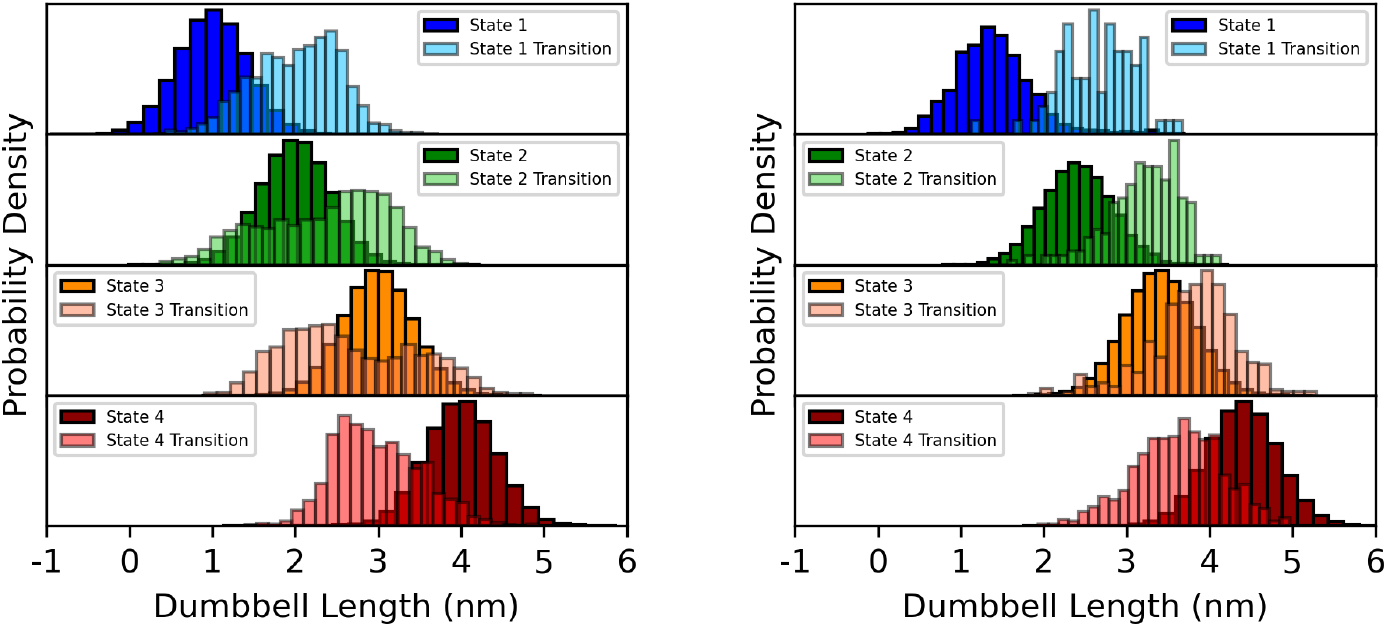
The normalised probability distributions, in the fully connected network system, of the dumbbell node separations and the separations specifically at the moment of transition into another mesostate. a) 0pN force applied. b) 10pN force applied.

The above is suggestive of alternative methods to parameterise the model from experiments. It may be the case that it is not possible to obtain *R_ij_* from equilibrium experimentation (e.g. if the rates are too slow, or too fast). A viable alternative, then, is to measure transition rates accelerated under the application of external forces. Such forces modify the energy landscape, but if these modifications are included in 14 and used to derive an alternate version of Eq.24, then the necessary 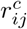 values can be obtained from the measured *R_ij_* under external force. Then, on removal of the force from the model, the equilibrium *R_ij_* (in the absence of force) may be inferred. This is a further example of how this mechano-kinetic framework can go beyond simply reproducing the initial parameterisation from experiment, extending the simulations to experimentally difficult and potentially infeasible scenarios for predictive purposes. The next section shows this formulation in the context of protein unfolding.

## Protein Unfolding

We now apply the framework to the problem of protein unfolding by mimicking SMFS-AFM experiments on polyproteins. At the same time, we will use protein unfolding as an example problem to investigate the differences between the two types of microstate model discussed in the Theoretical Development Section: the mechanical energy difference (MED) model and the force-activated transition (FAT) model. We note that conventional analysis for protein unfolding experiments is similar to the FAT model [32]: this makes sense as we might typically envisage that a force applied across a bond within the protein is what drives the unfolding. The kinetic schematic of this system, irrespective of model, is shown in Fig.2c, and the structural schematic is shown in Fig.8.

**Fig 8.**
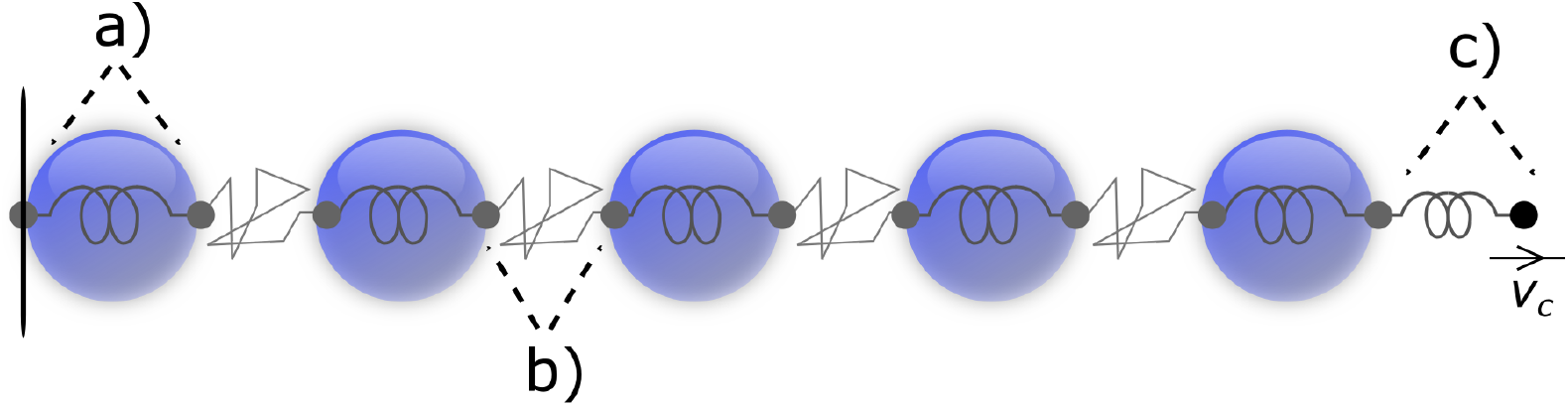
The structural model of our protein unfolding simulations for both the MED and FAT models. **a)** Each (one-dimensional) protein is modelled as a Hookean spring connecting two dynamic nodes. Upon a kinetic transition to the unfolded state, this spring transforms into a FENE spring, representing the unfolded amino-acid chain. Each node provides half the Stokes drag for each protein. The first node is frozen in place, as indicated by the vertical black line, to mimic the surface binding in SMFS-AFM experiments. **b)** The FENE spring representing the linker domain between protein subunits. **c)** Force is applied to the system by a bead attached to a spring, representing a cantilever. The bead moves at constant velocity and is independent of thermal noise, giving a well-characterised force-loading rate.

To model protein unfolding, we choose maltose binding protein (MBP) as a model protein due to the availability of all the required information to parameterise our simple 1D model. Rief *et al*. have studied MBP in great detail using SMFS-AFM [37] and further, equilibrium mechanical properties have been measured to a high accuracy using elastic incoherent neutron scattering (EINS) [38]. In addition, equilibrium unfolding behaviour has been experimentally inferred by Na *et al*. [17]. Finally, MBP has recently been used in cross-lengthscale mechanical studies of globular protein-based hydrogels [22], and so the model developed here may be extended to assist in the future understanding of unfolding in these systems.

We define two mesostates for each protein; one for the folded state and one for the unfolded. The folded state in a 1D model can be represented as a Hookean spring with an approximate spring constant *k*_1_ = 30.3pN/nm [38] and equilibrium length *l*_1_ = 5nm, approximately representing MBP [39]. The unfolded state can then be represented using a modified FENE potential, with associated force

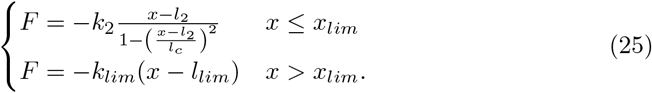

Eq.25 represents a non-interacting amino acid chain with only an entropic restoring force between the N- and C-termini at small extension. For *x* ≤ *x_lim_*, both the force and effective stiffness of the chain tend to infinity as *x* − *l*_2_ → *l_c_*, as the amino acid chain straightens and the large enthalpic contributions become apparent. *l_c_*, then, is the contour length of the polypeptide. This potential reproduces some of the core behaviour of the worm-like chain model, a more common model for polypeptides used in SMFS-AFM experimental analysis [32]. Unfortunately, the effective linear (Hookean) stiffness tending to infinity presents significant numerical problems, severely limiting the integration timestep we can take at large extensions. To compensate for this, we define the effective differential stiffness at each *x*, *k_eff_*, as

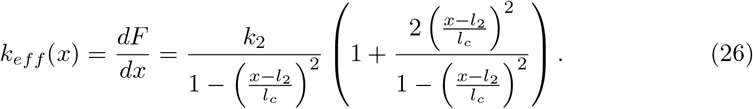

We relax the infinite barrier in the standard FENE potential by choosing a limiting stiffness *k_lim_*, such that when *k_eff_* > *k_lim_*, the potential becomes Hookean again with *k_lim_* being the new stiffness. This transition occurs at a value *x_lim_* (where *k_eff_* = *k_lim_*), and requires a new effective equilibrium length *l_lim_* chosen such that the force is continuous at *x* = *x_lim_*. Both *x_lim_* and *l_lim_* can be numerically calculated prior to the start of the simulation. We effectively use the FENE potential to transition from a linear stiffness *k*_2_ to a maximum linear stiffness *k_lim_*. We set *k_lim_* = 411pN/nm so that for our chosen *k_B_T* = 4.11pN.nm, fluctuations in the modified FENE chains are well-defined and negligible at large extension.

The average kinetic rates *R_ij_* in the absence of force are given in Table 7. The native unfolding rate of MBP was sourced from pulse proteolysis experiments [17], whereas the refolding rate was approximated using simple diffusion calculations i.e. the time taken for half the amino-acid chain to diffuse its own length. In practise, however, refolding almost never occurs in the simulation presented. These rates were used with Eq.14, and the relevant integrals were calculated numerically to account for the more complex modified FENE potential.

**Table 7.** The initial set of transition rates (in the absence of external force) used to investigate protein unfolding.

| Transition Rates, $R_{ij}$ ($\mu\text{s}^{-1}$) | | | $P_i$ |
| --- | --- | --- | --- |
| From / To | 1 | 2 |  |
| 1 | N/A | $6 \times 10^{-13}$ | $\sim 1$ |
| 2 | $2.15 \times 10^{-3}$ | N/A | $\sim 0$ |

As illustrated in Fig.8, we connect five proteins in series with shorter segments of modified FENE potential amino-acid chain with only 1 mesostate defined, representing the linker domain of an engineered polyprotein [32]. We use five primarily because relatively short polyproteins have previously been under independent investigation by the authors [40–43]. FENE parameters were calculated by using the limiting, soft case of the WLC model together with the number of amino acids in each chain, assuming that the persistence length of the chain *L_p_* = 0.4nm [44]. All parameters are given in Table 8.

**Table 8.**
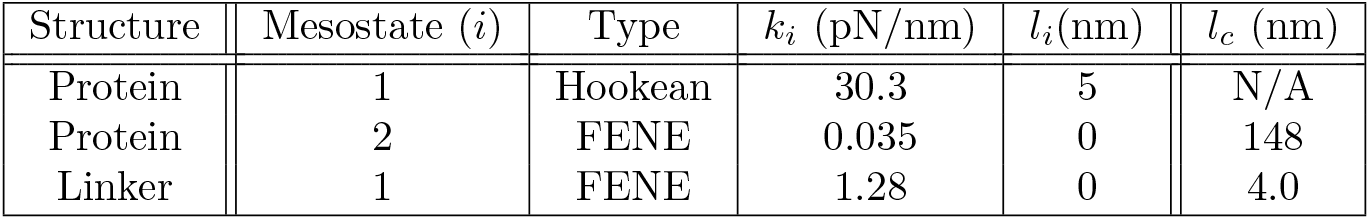
The energy function and material parameters used to define the protein mesostates used in the kinetic network of the polyprotein unfolding model.

In addition, the final protein in the series is connected to a linear spring, with spring constant *k_c_* = 1pN/nm, representing the cantilever of the atomic force microscope. This spring is in turn connected to a bead that is moved at a constant velocity *v_c_* in the +ve *x* direction. This progressively varies the force over the course of the simulation in a manner analogous to the stretching protocol employed in experiments [32, 40, 41], in which the base of the cantilever is moved with constant velocity, and identical to the constant-velocity steered MD protocol implemented in the NAMD MD package [45, 46]. All protein unfolding simulations were performed by also fixing the position of the left-most particle in the polyprotein. As such, for each particle *α* other than the first and last particles, each particle is subject to the a Langevin equation

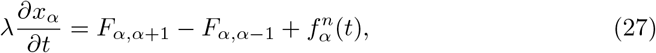

where *F_α,β_* is the elastic force applied to particle *α* by the connection to particle *β*, and is either a Hookean force (representing a folded protein), or a FENE force (representing either a linker or unfolded protein) depending on the spring object each pair of particles are connected by as the simulation progresses. The connecting object will change as part of the kinetic scheme. The initial particle is unique in that it never moves, and so isn’t subject to Langevin dynamics. The particle connected to the cantilever component is also unique, and has an extra force 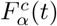 included instead of a further polyprotein component

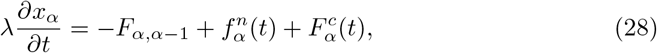

where *λ* = 6*π*pN.ns/nm and

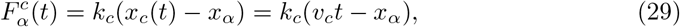

and leads to a well-characterised force-loading rate *R_L_* = *k_c_ v_c_* as the protein assembly reaches maximal extension (in practice the loading rate is slightly smaller due to the increasing extension of the proteins and linkers). We emphasise that the constant-velocity bead in our cantilever object is independent of thermal noise in our simulations, which is not the case experimentally.

Finally, when implementing the mechanical-energy difference model, we proceed as we have so far with the integrals in Eq.14 performed to determine each 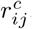. For the force-activated transition model, we require an additional parameter *b_ij_* for each integral in Eqs.17 and 18. In the first investigations we parametrise as best we can from available experimental data [37, 40], using *b_ij_* = 1nm as representative, but we later investigate the behaviour of this parameter as a part of the protein-unfolding simulation suite.

### Protein Unfolding - Results

We begin our investigation with an investigation of the effect of thermal noise on the measured unfolding force of the model proteins, implementing both the MED and FAT models. Thermal fluctuations, while small in comparison to work done by the externally applied force, are nevertheless included in the dynamic portion of the simulation, and therefore ought to directly affect the emergent kinetics. We set the stiffness of the constant-velocity spring component, *k_c_* = 1pN/nm, and the pulling speed, *v_c_* = 5 × 10^-4^nm/ns, to minimise the signal-noise ratio and so that the force increases at a reasonable rate. In the FAT implementation, we set *b_ij_* = *b* = 1.0nm for each protein along the chain, a somewhat arbitrary choice of unity but of the approximately the correct order of magnitude for protein unfolding [41] and representative of MBP itself [37].

Fig.9 shows four characteristic SMFS-AFM curves extracted from mechano-kinetic simulations performed with and without thermal noise for both model implementations. We emphasise, the force measured here on the y-axis is the external tension within the simulated cantilever components, not the internal force within any of the subunits. The extension on the x-axis is measured from the N-terminus of the first protein to the C-terminus of the last protein, not the position of the attached constant-velocity bead. This choice best represents what would be the tip of the cantilever in a real experiment.

**Fig 9.**
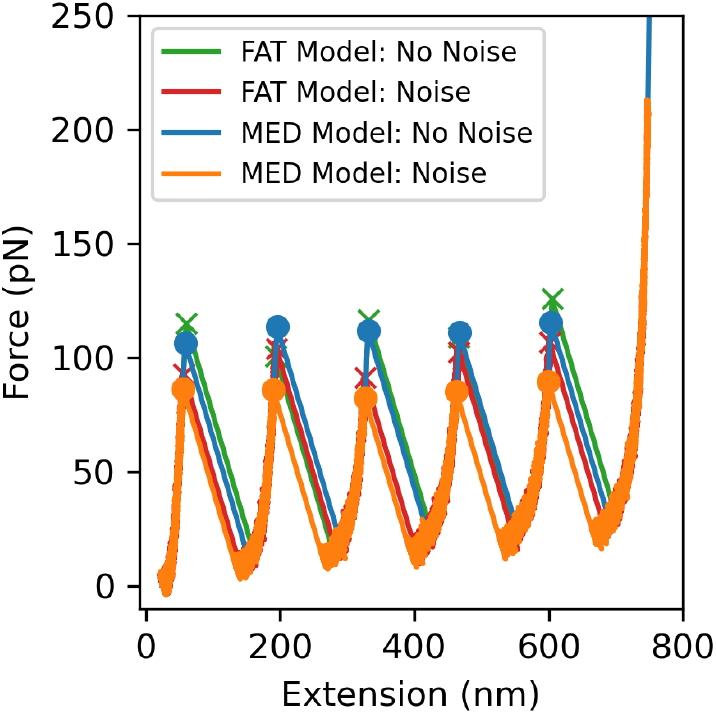
A set of polyprotein unfolding curves from associated protein unfolding simulations showing how the presence of thermal noise affects the unfolding kinetics.

We can see clearly that thermal noise has a significant effect on the unfolding force in both cases. By extracting the five peak forces (and their associated extensions), we can gain statistics for the unfolding kinetics. It has been previously observed that each specific peak has its own kinetic characteristics [47] but for now, we take the average of the these values from each simulation to gain single representative values for each system. For the MED model, in the absence of thermal noise, we measure an unfolding force, *F_un_* = 111.6 ± 1.4pN, and a peak-to-peak distance, *x_pp_* = 136.3 ± 0.3nm, and in the presence of thermal noise, we measure *F_un_* = 85.9 ± 1.0pN and *x_pp_* = 135.9 ± 0.7nm. For the FAT model, in the absence of thermal noise, we measure an unfolding force, *F_un_* = 113.5 ± 3.7pN, and a peak-to-peak distance, *x_pp_* = 136.3 ± 1.2nm, and in the presence of thermal noise, we measure *F_un_* = 99.5 ± 2.8pN and *x_pp_* = 136.3 ± 1.6nm. In all cases the measured *x_pp_* value is remarkably consistent, and the curves overlap well, implying physical consistency between simulations with and without noise. However, the measured *x_pp_* values are lower than *l_c_* − (*l*_1_ + *F_un_*/*k*_1_), which is the value we would expect if the FENE chain were to fully extend under the externally applied force. This implies that the proteins gain enough energy to unfold at slightly lower extensions. Nevertheless, this difference is relatively small. More importantly, the presence of thermal noise lowers the required unfolding force by approximately 25pN in the MED model, and 14pN in the FAT model. This difference in the reduction in *F_un_* upon the inclusion of thermal noise is outside the uncertainty, suggesting a real difference between the models, but may be due to the difference in the value of *b* used and the average distance to the unfolding transition in the MED model. In both cases however, such a reduction in *F_un_* was quite unexpected as thermal fluctuations contribute a modest 0.5*k_B_T* of mechanical energy on average at zero external force, a negligible impact on the stability of most proteins, and MBP in particular. On reflection, however, due to the exponential nature of Boltzmann’s equation, we see that the same amount of energy at high force has a relatively large impact, pushing the protein over the energy barrier required to unfold. This is a significant validation of the relevance of thermal noise in biological systems beyond being simply a tool for diffusion and entropic exploration of conformational space. This will be the case especially in non-equilibrium simulations and in models of proteins with low barriers to unfolding, such as intrinsically disordered proteins [48] and some extremophilic proteins [49–51]. All simulations from this point forward will be performed with thermal noise included.

We now investigate the effect on this system of variations in the (folded) protein spring constant *k*_1_, and the pulling speed *v_c_*. It is well known that changes in the pulling speed ought to alter the measured unfolding force due to the unfolding mechanism itself, but interestingly, West *et al*. also predicted that the existence of internal dynamics ought to alter the unfolding transition kinetics [26], and varying k1 ought to allow a systematic analysis of this. First using the MED model, we performed a single unfolding simulation for each of the parameter pairs defined in Table 9, and again extracted the unfolding forces, *F_un_*, and and peak-to-peak distances, xpp. All other parameters were kept constant between simulations, including the average rates *R_ij_*.

**Table 9.** The parameter space investigated to determine the effect on the unfolding forces and peak-to-peak separations of protein unfolding curves.

| Parameter | Values |  |  |  |
| --- | --- | --- | --- | --- |
| $k_1$ (pN/nm) | 10.0 | 30.3 | 50.0 | |
| $v_c$ (nm/ns) | $5 \times 10^{-4}$ | $1 \times 10^{-3}$ | $5 \times 10^{-3}$ | $1 \times 10^{-2}$ |

From Fig.10, in the MED model we observe that the protein stiffness has the most significant effect on both *F_un_* for all values of *v_c_* (for a given value of *R_ij_*). Rewriting the linear protein energy explicitly in terms of force, 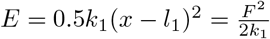, we see that for a higher stiffness, a higher force is required to reach the same energy. Thus, we see that the unfolding force increases with the internal stiffness of the protein. Interestingly, the peak-to-peak separation *x_pp_* also increases with internal protein stiffness. As a function of the cantilever pulling speed *v_c_*, however, we see too much variation between peak-to-peak values to draw any conclusions. However, it is clear that the unfolding force does slightly increase with *v_c_*, as expected.

**Fig 10.**
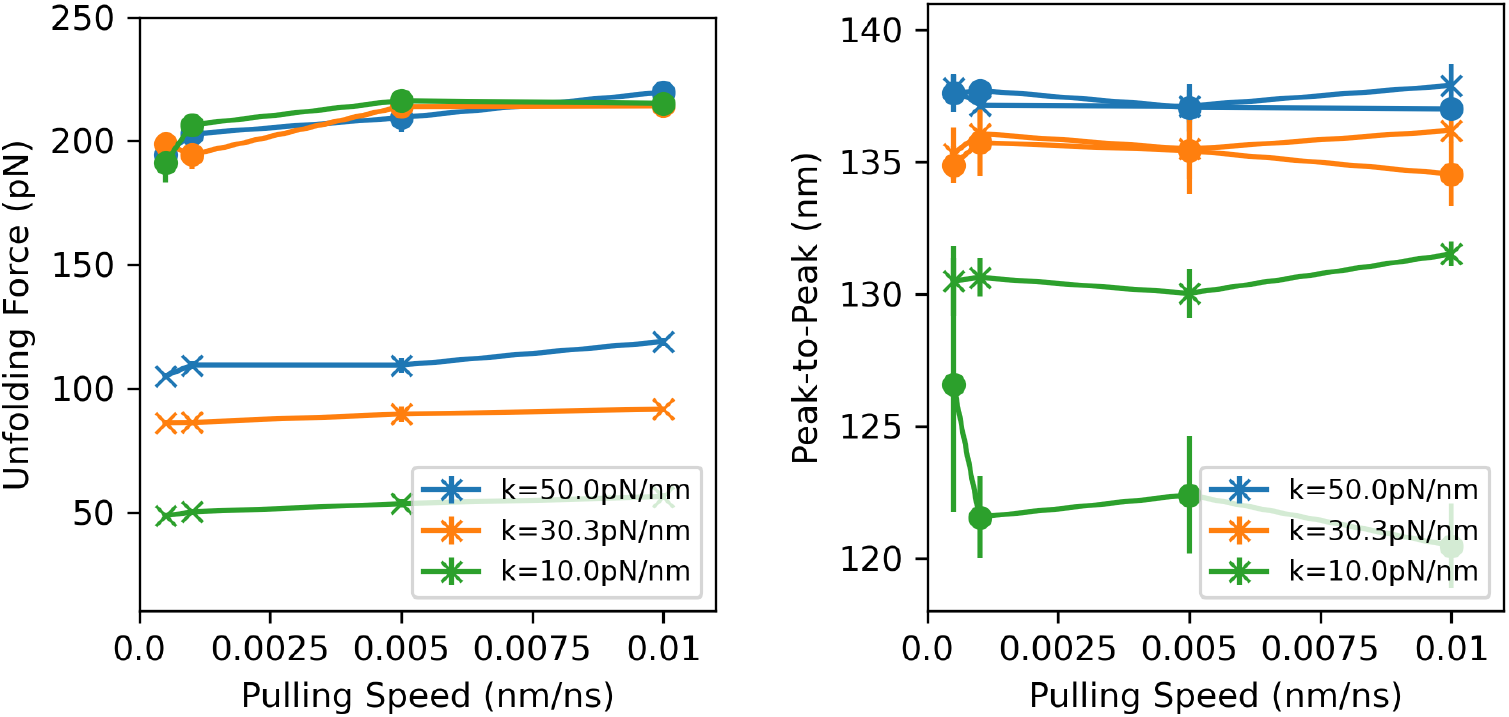
A series of protein unfolding simulations using both the MED model (crosses) and the FAT model (circles) showing how the cantilever pulling speed and protein stiffness affects the unfolding kinetics. a) Unfolding forces. b) Peak-to-peak distances.

We performed an identical set of simulations within the FAT model, setting the parameter *b* = 0.5nm, again of the correct order of magnitude for (MBP) protein unfolding [37, 41], but also to more clearly show the difference between these models in the parameter sweep. We again see that the peak-to-peak separations are most dominated by the protein stiffness, with the most pronounced difference from the MED model occurring for *k*_1_ = 10pN/nm. We also see that the unfolding forces are much greater than those in the MED model in each case. This will later be shown to be a simple matter of effective parameterisation differences between models. More importantly, though, we observe the unfolding forces in each case are, as expected from Eq.15, approximately independent of protein stiffness. The two models are therefore clearly unique, showing that our choice of mechanical model in the framework is inseparable from the results we obtain. Thus, in the absence of knowledge of the “true” underlying mechanical model, the full mechano-kinetic framework acts as a tool to investigate what we may expect to see, given a certain set of assumptions and experimental data.

To emphasis this point, we will now use the protein unfolding simulations to investigate two parameters unique to each model. First, we observe the effect of the set of *f_ij_* in the MED model. These values were (to our knowledge) first used conceptually by Sarlah *et al*. to imply a “mechanical activation energy”, and modify how far towards the activation energy we need to be to perform the discretised kinetic transition [23]. A value of *f_ij_* close to 1 implies that the mechanical energy barrier is close to the target state, and a value close to zero implies that the mechanical energy barrier is close to the initial state. Fig.11 shows the results of altering *f_ij_* on the protein unfolding simulations with *k*_1_ = 30.3 pN/nm and *v_c_* = 10^-3^nm/ns, where *f*_01_ on the x-axis represents the value used for the unfolding transition.

**Fig 11.**
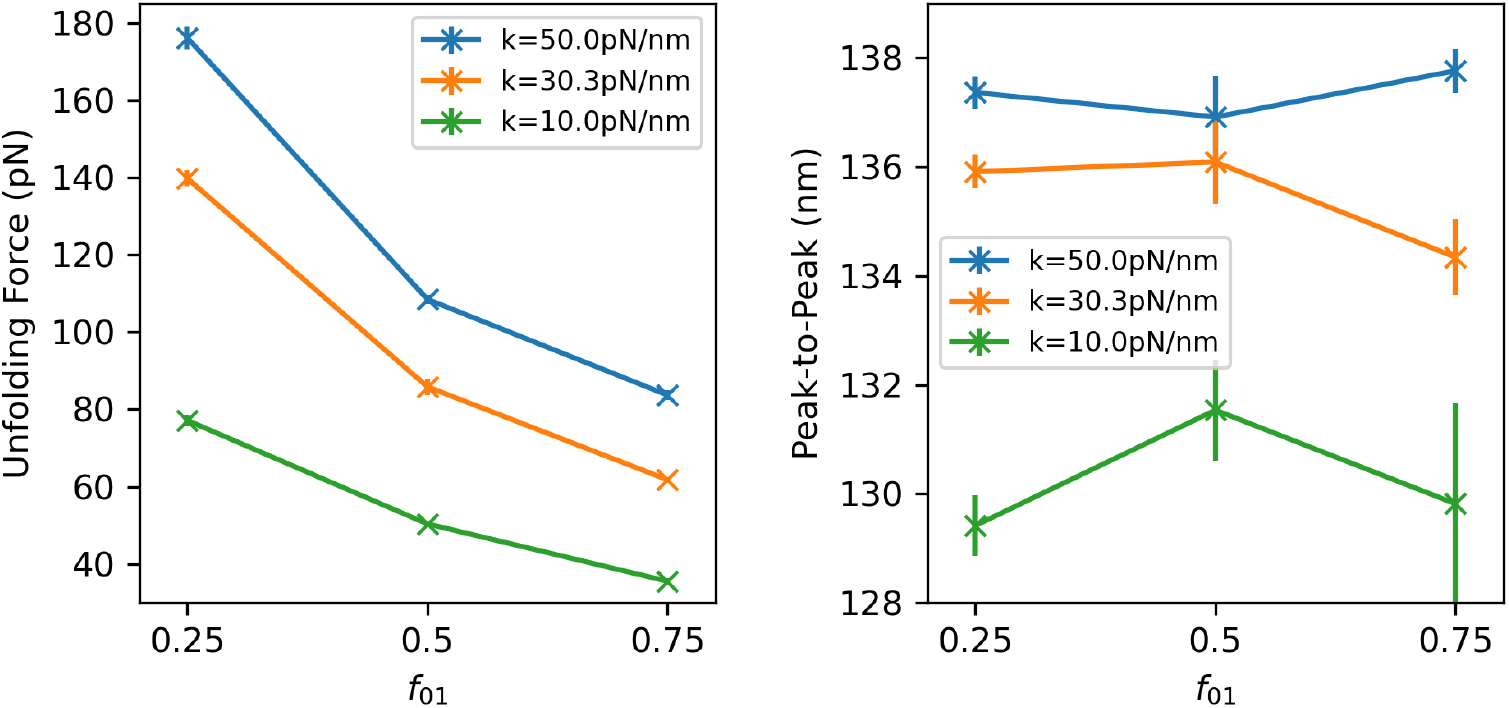
A series of protein unfolding simulations, varying *f_ij_* and *k*_1_ a) Unfolding forces. b) Peak-to-peak distances.

We see that the effect of varying *k*_1_ on both *F_un_* and *x_pp_* is qualitatively the same across all values of *f*_01_, increasing both the unfolding force and peak-to-peak distance as previously observed, showing that the *f_ij_* parameters do act as scaling factors with respect to the energy term. Again, *x_pp_* is not much affected by the variations in *f*_01_, but the unfolding force *F_un_* shows a significant decrease. It is intuitive to see that, because the multiplication of *f_ij_* and 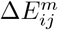 together are what determine the transition rate, an increase in *f*_01_ would mean that a smaller 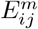 is required for the same transition probability, and thus a lower unfolding force is observed. However, a more interesting perspective emerges from the observation that *f_ij_* relates to something like a mechanical activation energy. As 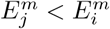, then increasing *f_ij_* reduces the effective mechanical activation energy required to make the transition, thus reducing the unfolding force.

To conclude this investigation, we now investigate the effect of the value *b_ij_* on the FAT model. In this model, we recall that the protein internal force is used continuously to modulate the kinetic transition rates, together with a distance parameter *b_ij_*. *b_ij_* is a parameter we set in advance, and so we again set *b_ij_* = *b* the same for all proteins along the chain. We set *k*_1_ = 30.3 pN/nm and performed a series of simulations for various values of *b* and *v_c_*. As each protein unfolded, we recorded the unfolding force within the cantilever object (as would be measured experimentally), 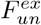, the unfolding force within the protein itself, *f_un_*(*x*) and therefore also the specific mechanical extension of the protein, Δ*x*.

In standard theoretical interpretations of protein unfolding experiments, the Bell-Evans-Ritchie model is used to calculate the effective unfolding rates as a function of external force, *R_ij_*(*F^ex^*) [32]:

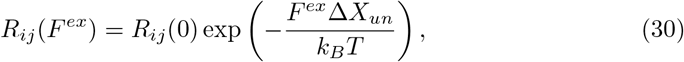

where *F^ex^* is some externally applied force and Δ*X_un_* is the “distance” through the free-energy landscape to the transition barrier. *R_ij_* (0), then, is the same *R_ij_* we have been using throughout this work, the experimentally measured equilibrium unfolding rate at the mesoscale. This equation is similar in form to Eq.15, but with important differences. First, although the value *b* is included at the microscale, we first show that it is equivalent to value Δ*X_un_* used in Eq.30. From the work of Evans [52], Hughes *et al*. [32] were able to show that the expected unfolding force as a function of relevant variables, 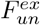, is:

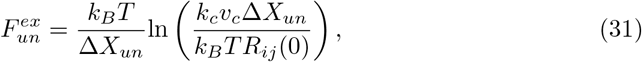

Using *b* = Δ*X_un_* we fit Eq.31 to our simulation data, shown in Fig.12a). We find good agreement between our simulations and theoretical predictions, indicating that *b* is indeed likely equivalent to Δ*X_un_*. This is shown more explicitly in Fig.12b), where we have plotted a characteristic energy *E_ch_* for both the theoretical predictions and our measured results. By removing the *b* dependence for the RHS of Eq.31, we define:

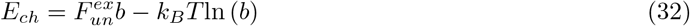

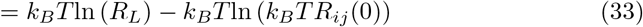

where *R_L_* = *k_c_ v_c_* is the force-loading rate. We enforce a gradient of *k_B_T* in our linear fit of Eq.32 to the data, thus implicitly assuming that any errors are due to differences in the effective transition rates, and find an approximate difference of 6pN.nm between our results and the theoretically expected values. This is a minor deviation on the order of thermal fluctuations at equilibrium. The *E_ch_* values in the Fig.12b) legend correspond to extrapolations to the experimentally used loading rates, assuming *v_c_* = 6 × 10^-7^nm/ns and *k_c_* = 50pN/nm. Interestingly, Hoffmann *et al*. also calculated a related form of characteristic energy in their study of many naturally occurring proteins with differing measured values of Δ*X_un_* and *F_un_*. From this collection they determined a typical *E_ch_* = 39.4 ± 3.1pN.nm for each protein in their survey. However, in an earlier work [40], they also found MBP to be a protein which significantly differs from all others, suggesting that the initial unfolding rate Rij may be responsible for this huge difference between MBP and other natural proteins.

**Fig 12.**
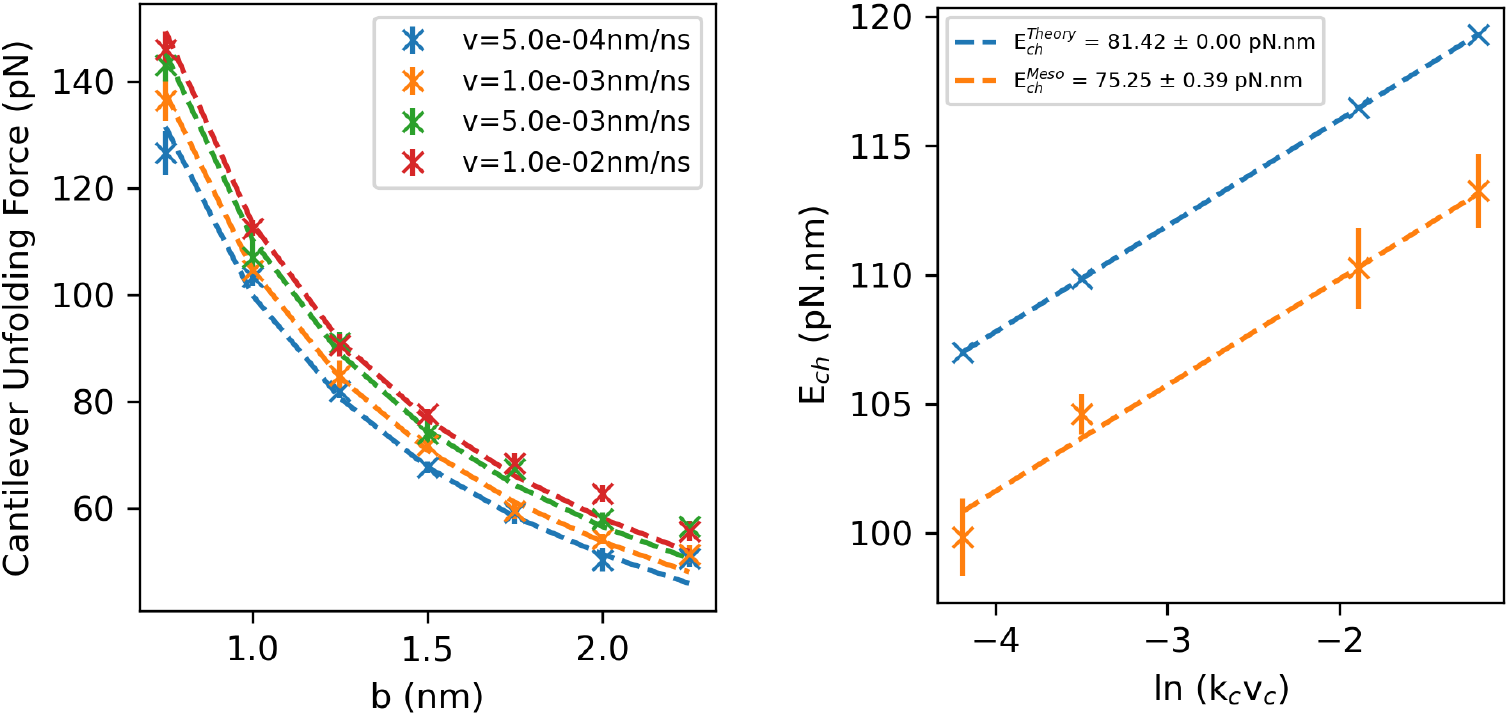
A series of protein unfolding simulations, varying *b*, for a variety of cantilever speeds vc. We observe the approximately correct theoretical dependence of *b* and the unfolding force, 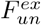. a) Unfolding force traces. b) Fit and extrapolation of characteristic unfolding energies *E_ch_* to the experimental loading rates *R_L_* = *k_c_ v_c_*.

It is clear that the FAT model is likely a more appropriate model for the problem of protein unfolding than the MED model, given the success of reproduction of existing theoretical predictions.

## Discussion

We have developed a “mechano-kinetic” simulation framework which enables instantaneous kinetic transitions to occur within, and be explicitly affected by, an underlying dynamic simulation framework. By integrating out the desired mechanical contributions to the (experimentally measured) free energies, expressed as transition rates between discrete kinetic states, we can infer the dynamics present at the microscale for a given energy landscape. We applied this framework to a simple dumbbell simulation and showed that our model correctly reproduces the free energy landscape used to parametrise the model, while at the same time showing exactly where in phase space kinetic transitions are likely to occur, again given a set of experimentally observed mesostate and an appropriate mechanical model. We further showed that external potentials can be introduces to the model, and found that under the action of external forces, while the connectivity of the kinetic states do not affect the measured mesostate occupation probabilities, they strongly affect the microstate populations and, importantly, the distribution of transition microstates. We finally applied the framework to the case of protein unfolding, and showed that characteristic SFMS-AFM unfolding traces can be produced. Utilising both the “Mechanical energy difference” model and the “Force-activated transition” model, we observe that our selection of the underlying mechanical energy landscape and state transition model are inseparable from the interpretations of the results of any simulations.

The original intention of this project was to develop an empirical framework, enabling extremely complex biophysical events (i.e. protein unfolding, conformational changes, binding / unbinding of ligands etc) to be implicitly simulated in parallel with a dynamic model without the need for extremely high resolution and the large associated amounts of computing power. We have achieved this goal, and even though our protein unfolding simulations show slight deviation from theoretical predictions, we can still calibrate the model to give the behaviour we desire.

Going forward, we note that there also exist systems in which the energy barrier between kinetic states is naturally quite low as compared to typical thermal energies [48–51]; in such cases it is difficult to make a clear distinction between states because transitions between them happen readily and frequently. Intermediate states have been observed in our example systems [37, 53]. The development of Markov state models (MSMs) [54, 55] shows that discrete kinetic states can be defined somewhat arbitrarily, and as such energy barriers can be arbitrarily low. Finally, Bialek *et al*. noted early on that protein dynamics and their reaction rates are coupled [56], such that a majority of the reactive events in some proteins can be coupled to specific protein modes of motion, indicating dynamic specificity. Multiple groups discovered almost in parallel that SMFS-AFM yields different unfolding forces depending on the directions in which you apply force, indicating dynamic anisotropy [57–59]. In these cases, then, the instantaneous location in physical space and associated mechanical energies can have a non-negligible effect on the overall free energies, and thus the transition rates between states. The inspiring work of Trott *et al*. emphasises this, introducing a highly detailed mechanistic model of cytoplasmic dynein which combines a wealth of experimental evidence to deduce the continuous function of the motor [60]. Examination of their computational insights indicates that from an atomistic statistical mechanics perspective, the mesoscopic transition rates emerge from the underlying dynamical behaviour.

Our model can account for all of these possibilities. With an appropriate definition of a mechanical energy landscape, whether high-resolution molecular dynamics or bespoke, including velocity degrees of freedom if necessary, mesostate definitions and the transition mechanisms between them, new insights can be drawn into the coupling between microstate dynamic behaviour and mesoscale kinetic behaviour in systems traditionally thought to be too far separated in characteristic timescale to simulate both in parallel.

## Supporting information

Supporting Information and Verification of Results

## Acknowledgments

We would like to thank Matt D.G. Hughes, Christa Brown, Harrison Laurent, Kalila Cook and Mazin Nasralla for discussions of the model in the context of protein unfolding in complex systems. Ben Hanson was initially supported by the EPSRC through a DTA, and completed the work supported again by the EPSRC with grant EP/P02288X/1 awarded to Professor Lorna Dougan.

## References

1. Bates M, Huang B, Rust MJ, Dempsey GT, Wang W, Zhuang X. Sub-Diffraction-Limit Imaging with Stochastic Optical Reconstruction Microscopy. In: A G, R R, J W, editors. Single Molecule Spectroscopy in Chemistry, Physics and Biology. Springer Berlin Heidelberg; 2010.

2. Cheng Y. Single-particle Cryo-EM at crystallographic resolution. Cell. 2015;161(3):450–457.

3. Kühlbrandt W. The Resolution Revolution. Science. 2014;343(6178):1443–1444.

4. Frank J. Time-resolved Cryo-Electron Microscopy: Recent Progress. Journal of Structural Biology. 2017;200(3):303–306.

5. Schermelleh L, Ferrand A, Huser T, Eggeling C, Sauer M, Biehlmaier O, et al. Super-resolution microscopy demystified. Nature Cell Biology. 2019;21(1):72–84.

6. Lawson CL, Baker ML, Best C, Bi C, Dougherty M, Feng P, et al. EMDataBank.org: Unified data resource for CryoEM. Nucleic Acids Research. 2011;39:456–464.

7. Solernou A, Hanson BS, Richardson RA, Welch R, Read DJ, Harlen OG, et al. Fluctuating Finite Element Analysis (FFEA): A continuum mechanics software tool for mesoscale simulation of biomolecules. PLoS Computational Biology. 2018;14(3):1–6.

8. Leng J, Shoura M, McLeish TCB, Real AN, Hardey M, McCafferty J, et al. Securing the future of research computing in the biosciences. PLoS Computational Biology. 2019;15(5):1–15.

9. Richardson RA, Hanson BS, Read DJ, Harlen OG, Harris SA. Exploring the dynamics of flagellar dynein within the axoneme with Fluctuating Finite Element Analysis. Quarterly Reviews of Biophysics. 2020;53.

10. Hanson BS, Iida S, Read DJ, Harlen OG, Kurisu G, Nakamura H, et al. Continuum mechanical parameterisation of cytoplasmic dynein from atomistic simulation. Methods. 2021;185:39–48.

11. Kubo S, Li W, Takada S. Allosteric conformational change cascade in cytoplasmic dynein revealed by structure-based molecular simulations. PLoS Computational Biology. 2017;13(9):1–27.

12. Reck-peterson SL, Yildiz A, Carter AP, Gennerich A, Vale RD, Mcintosh JR, et al. Single-Molecule Analysis of Dynein Processivity and Stepping Behavior. Cell. 2006;126(2):335–348.

13. Li L, Alper J, Alexov E. Cytoplasmic dynein binding, run length, and velocity are guided by long-range electrostatic interactions. Scientific Reports. 2016;6(1):1–12.

14. Kinoshita Y, Kambara T, Nishikawa K, Kaya M, Higuchi H. Step Sizes and Rate Constants of Single-headed Cytoplasmic Dynein Measured with Optical Tweezers. Scientific Reports. 2018;8(1):1–14.

15. Sasaki K, Kaya M, Higuchi H. A Unified Walking Model for Dimeric Motor Proteins. Biophysical Journal. 2018;115(10):1981–1992.

16. Dudko OK, Hummer G, Szabo A. Intrinsic rates and activation free energies from single-molecule pulling experiments. Physical Review Letters. 2006;96(10):1–4.

17. Na YR, Park C. Investigating protein unfolding kinetics by pulse proteolysis. Protein Science. 2009;18(2):268–276.

18. West DK, Olmsted PD, Paci E. Mechanical unfolding revisited through a simple but realistic model. Journal of Chemical Physics. 2006;124(15).

19. Ohashi KG, Han L, Mentley B, Wang J, Fricks J, Hancock WO. Load-dependent detachment kinetics plays a key role in bidirectional cargo transport by kinesin and dynein. Traffic. 2019;20(4):284–294.

20. Belyy V, Schlager MA, Foster H, Reimer AE, Carter AP, Yildiz A. The mammalian dynein-dynactin complex is a strong opponent to kinesin in a tug-of-war competition. Nature Cell Biology. 2016;18(9):1018–1024.

21. Linke WA, Ivemeyer M, Mundel P, Stockmeier MR, Kolmerer B. Nature of PEVK-titin elasticity in skeletal muscle. Proceedings of the National Academy of Sciences of the United States of America. 1998;95(14):8052–8057.

22. Hughes MDG, Cussons S, Mahmoudi N, Brockwell DJ, Dougan L. Single molecule protein stabilisation translates to macromolecular mechanics of a protein network. Soft Matter. 2020;16(27):6389–6399.

23. Sarlah A, Vilfan A. The winch model can explain both coordinated and uncoordinated stepping of cytoplasmic dynein. Biophysical Journal. 2014;107(3):662–671.

24. Sarlah A, Vilfan A. Minimum requirements for motility of a processive motor protein. PLoS ONE. 2017;12(10):1–13.

25. West DK, Olmsted PD, Paci E. Free energy for protein folding from nonequilibrium simulations using the Jarzynski equality. Journal of Chemical Physics. 2006;125(20):1–7.

26. West DK, Paci E, Olmsted PD. Internal protein dynamics shifts the distance to the mechanical transition state. Physical Review E. 2006;74(6):061912.

27. Staple DB, Payne SH, Reddin ALC, Kreuzer HJ. Model for stretching and unfolding the giant multidomain muscle protein using single-molecule force spectroscopy. Physical Review Letters. 2008;101(24):1–4.

28. Sikora M, Sulkowska JI, Witkowski BS, Cieplak M. BSDB: The biomolecule stretching database. Nucleic Acids Research. 2011;39(SUPPL. 1).

29. Booth JJ, Shalashilin DV. Fully Atomistic Simulations of Protein Unfolding in Low Speed Atomic Force Microscope and Force Clamp Experiments with the Help of Boxed Molecular Dynamics. Journal of Physical Chemistry B. 2016;120(4):700–708.

30. Takada S. Go model revisited. Biophysics and Physicobiology. 2019;16(0):248–255.

31. Bell GI. Models for the specific adhesion of cells to cells. Science. 1978;200(4342):618–627.

32. Hughes ML, Dougan L. The physics of pulling polyproteins: A review of single molecule force spectroscopy using the AFM to study protein unfolding. Reports on Progress in Physics. 2016;79(7):076601.

33. Hughes MDG, Hanson BS, Cussons S, Mahmoudi N, Brockwell DJ, Dougan L. Control of Nanoscale in Situ Protein Unfolding Defines Network Architecture and Mechanics of Protein Hydrogels. ACS Nano. 2021;15(7):11296–11308.

34. Imai H, Shima T, Sutoh K, Walker ML, Knight PJ, Kon T, et al. Direct observation shows superposition and large scale flexibility within cytoplasmic dynein motors moving along microtubules. Nature Communications. 2015;6(1):1–11.

35. Kamiya N, Mashimo T, Takano Y, Kon T, Kurisu G, Nakamura H. Elastic properties of dynein motor domain obtained from all-atom molecular dynamics simulations. Protein Engineering, Design and Selection. 2016;29(8):317–326.

36. Niekamp S, Stuurman N, Zhang N, Vale RD. Three-color single-molecule imaging reveals conformational dynamics of dynein undergoing motility. Proceedings of the National Academy of Sciences of the United States of America. 2021;118(31).

37. Bertz M, Rief M. Mechanical Unfoldons as Building Blocks of Maltose-binding Protein. Journal of Molecular Biology. 2008;378(2):447–458.

38. Wood K, Frölich A, Paciaroni A, Moulin M, Härtlein M, Zaccai G, et al. Coincidence of dynamical transitions in a soluble protein and its hydration water: Direct measurements by neutron scattering and MD simulations. Journal of the American Chemical Society. 2008;130(14):4586–4587.

39. Telmer PG, Shilton BH. Insights into the Conformational Equilibria of Maltose-binding Protein by Analysis of High Affinity Mutants. Journal of Biological Chemistry. 2003;278(36):34555–34567.

40. Hoffmann T, Dougan L. Single molecule force spectroscopy using polyproteins. Chemical Society Reviews. 2012;41(14):4781–4796.

41. Hoffmann T, Tych KM, Hughes ML, Brockwell DJ, Dougan L. Towards design principles for determining the mechanical stability of proteins. Physical Chemistry Chemical Physics. 2013;15(38):15767–15780.

42. Tych KM, Hughes ML, Bourke J, Taniguchi Y, Kawakami M, Brockwell DJ, et al. Optimizing the calculation of energy landscape parameters from single-molecule protein unfolding experiments. Physical Review E - Statistical, Nonlinear, and Soft Matter Physics. 2015;91(1):1–9.

43. Da Silva MA, Lenton S, Hughes M, Brockwell DJ, Dougan L. Assessing the Potential of Folded Globular Polyproteins As Hydrogel Building Blocks. Biomacromolecules. 2017;18(2):636–646.

44. Stirnemann G, Giganti D, Fernandez JM, Berne BJ. Elasticity, structure, and relaxation of extended proteins under force. Proceedings of the National Academy of Sciences of the United States of America. 2013;110(10):3847–3852.

45. Phillips JC, Hardy DJ, Maia JDC, Stone JE, Ribeiro JV, Bernardi RC, et al. Scalable molecular dynamics on CPU and GPU architectures with NAMD. Journal of Chemical Physics. 2020;153(4).

46. Mücksch C, Urbassek HM. Accelerating Steered Molecular Dynamics: Toward Smaller Velocities in Forced Unfolding Simulations. Journal of Chemical Theory and Computation. 2016;12(3):1380–1384.

47. Zinober RC, Brockwell DJ, Beddard GS, Blake AW, Olmsted PD, Radford SE, et al. Mechanically unfolding proteins: The effect of unfolding history and the supramolecular scaffold. Protein Science. 2009;11(12):2759–2765.

48. Uversky VN. Intrinsically disordered proteins and their ‘‘Mysterious” (meta)physics. Frontiers in Physics. 2019;7:8–23.

49. Feller G. Protein stability and enzyme activity at extreme biological temperatures. Journal of Physics Condensed Matter. 2010;22(32).

50. Reed CJ, Lewis H, Trejo E, Winston V, Evilia C. Protein adaptations in archaeal extremophiles. Archaea. 2013;.

51. Brininger C, Spradlin S, Cobani L, Evilia C. The more adaptive to change, the more likely you are to survive: Protein adaptation in extremophiles. Seminars in Cell and Developmental Biology. 2018;84:158–169.

52. Evans E. Probing the Relation Between Force, Lifetime and Chemistry in Single Molecular Bonds. Annual Review of Biophysics and Biomolecular Structure. 2001;30(1):105–128.

53. Isojima H, Iino R, Niitani Y, Noji H, Tomishige M. Direct observation of intermediate states during the stepping motion of kinesin-1. Nature Chemical Biology. 2016;12(4):290–297.

54. Bowman GR, Pande VS, Noe F. An Introduction to Markov State Models and Their Application to Long Timescale Molecular Simulation. Springer; 2014.

55. Husic BE, Pande VS. Markov State Models: From an Art to a Science. Journal of the American Chemical Society. 2018;140(7):2386–2396.

56. Bialek W, Onuchic JN. Protein dynamics and reaction rates: mode-specific chemistry in large molecules? Proceedings of the National Academy of Sciences of the United States of America. 1988;85(16):5908–5912.

57. Brockwell DJ, Paci E, Zinober RC, Beddard GS, Olmsted PD, Smith DA, et al. Pulling geometry defines the mechanical resistance of a beta-sheet protein. Nature Structural Biology. 2003;10(9):731–737.

58. Carrion-Vazquez M, Li H, Lu H, Marszalek PE, Oberhauser AF, Fernandez JM. The mechanical stability of ubiquitin is linkage dependent. Nature Structural Biology. 2003;10(9):738–743.

59. Dietz H, Berkemeier F, Bertz M, Rief M. Anisotropic deformation response of single protein molecules. Proceedings of the National Academy of Sciences of the United States of America. 2006;103(34):12724–12728.

60. Trott LE, Hafezparast M, Madzvamuse A. A mathematical understanding of how cytoplasmic dynein walks on microtubules. Royal Society Open Science. 2018;5(8):171568.

