## Supporting Information and Verification of Results for "A generalised mechano-kinetic model for use in multiscale simulation protocols"

### Data Access & Reproducibility

All of our simulations were performed using bespoke protocols designed specifically for purpose. The software is available for download from the Bitbucket repository `https://bitbucket.org/GokuBH/1dmechanokineticmodel/`. Although the core software may be modified for future use, for reproducibility purposes the branch *Publication* remains unchanged since this work was performed.

All data and graphs used in this work, as well as additional movies which comprise the range of input parameterisations used in this work, will be made available online upon publication.

### Validation of Mechano-Kinetic Simulations

In all of the following simulations, as in the main article, we will set the drag, $\lambda = 6\pi$pN.ns/nm for each sphere, and temperature $T = 298$K, giving a thermal energy $k_B T \approx 4.11$ pN.nm.

#### Dynamic Model Validation

We first validate our implementation of the dynamic dumbbell model by running a simulation with a single kinetic state i.e. with no kinetic transitions. We set $k = 10$pN/nm and $l = 0$nm. Fig.S1 shows the convergence of physically measurable properties to their analytical predictions. Fig.S1a) shows that the single degree of freedom converges to $\langle U \rangle = {}^1\!/_2 k_B T$ as expected, indicating that our transition rates are sufficiently low to correspond to equilibrium statistical mechanics within each of the mesostates. Fig.S1b) shows the centroid of the system diffusing as a single object with a viscous drag equal to the sum of the two drags of the individual nodes, indicating correct diffusional behaviour and that the internal elastic forces are conservative.

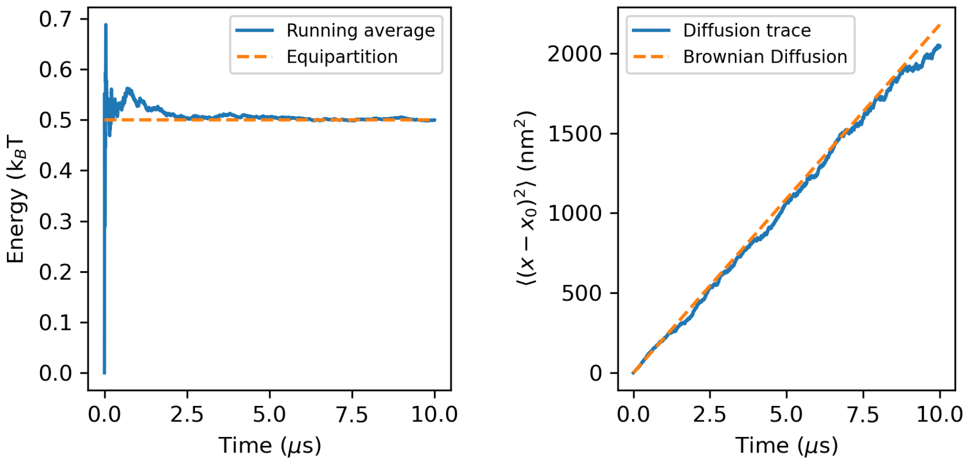


**Fig S1.** The physical characteristics emerging from our dynamical model. **a)** Elastic energy, calculated as an average evolving over the course of a single simulation. **b)** Diffusion, where the positional variance is calculated as an average over 400 repeat simulations.

| Mesostate ($i$) | $k_i$ (pN/nm) | $l_i$(nm) |
|---|---|---|
| 1 | 1 | 4 |
| 2 | 2 | 3 |
| 3 | 3 | 2 |
| 4 | 4 | 1 |

**Table S1.** The material parameters used to define the mesostates in a fully connected kinetic network of the dumbbell model.

| Transition Rates, $R_{ij}$ ($\mu s^{-1}$) | | | | | $P_i$ |
|---|---|---|---|---|---|
| From / To | 1 | 2 | 3 | 4 | |
| 1 | N/A | 0.3 | 0.2 | 0.1 | 0.1 |
| 2 | 0.4 | N/A | 0.2 | 0.1 | 0.2 |
| 3 | 0.4 | 0.3 | N/A | 0.1 | 0.3 |
| 4 | 0.4 | 0.3 | 0.2 | N/A | 0.4 |

| Transition Rates, $R_{ij}$ ($\mu s^{-1}$) | | | | | $P_i$ |
|---|---|---|---|---|---|
| From / To | 1 | 2 | 3 | 4 | |
| 1 | N/A | 3 | 2 | 1 | 0.1 |
| 2 | 4 | N/A | 2 | 1 | 0.2 |
| 3 | 4 | 3 | N/A | 1 | 0.3 |
| 4 | 4 | 3 | 2 | N/A | 0.4 |

**Table S2.** Two matrices defining sets of average transition rates between mesostates, $R_{ij}$, used in a fully connected kinetic network of the dumbbell model, together with the associated mesostate occupation probabilities calculated using the principle of detailed balance.

### Kinetic Model Validation

To validate our implementation of the kinetic dumbbell model we consider four different parametrisations of the dumbbell model as detailed in Table S1. These correspond to four different mesostates, each with a different local energy landscape. This kinetic landscape was designed to have the resultant property (if successful) of an equally distributed, inhomogeneous mesostate probability distribution for testing purposes. As per detailed balance we can attain this probability distribution with either of the sets of average transition rates, $R_{ij}$, shown in Table S2. The second set of rates is (uniformly) an order of magnitude faster. As there are no off-diagonal zero rates contained within these transition rate matrices, this model corresponds to a fully connected kinetic network.

We calculated the evolution of the occupation probabilities for each mesostate within a single simulation, shown in Fig.S2. We observe convergence to the expected mesostate occupation probabilities, given the set of average rates in Table S2 which were used by the simulation (possibly representing experimentally measured rates of a more realistic system). We see that the simulation in in Fig.S2a), corresponding to the lower set of rates, converges approximately ten times slower than the simulation in Fig.S2b), indicating that our simulation is working correctly at the mesostate level. We emphasise that this convergence was achieved whilst each individual transition between microstates in different mesostates, $(\vec{x})_i$ and $(\vec{x})_j$, was explicitly dependent upon the current mechanical energy of the system, showing our generalised coupling between microstate dynamics and mesostate kinetics to be valid.

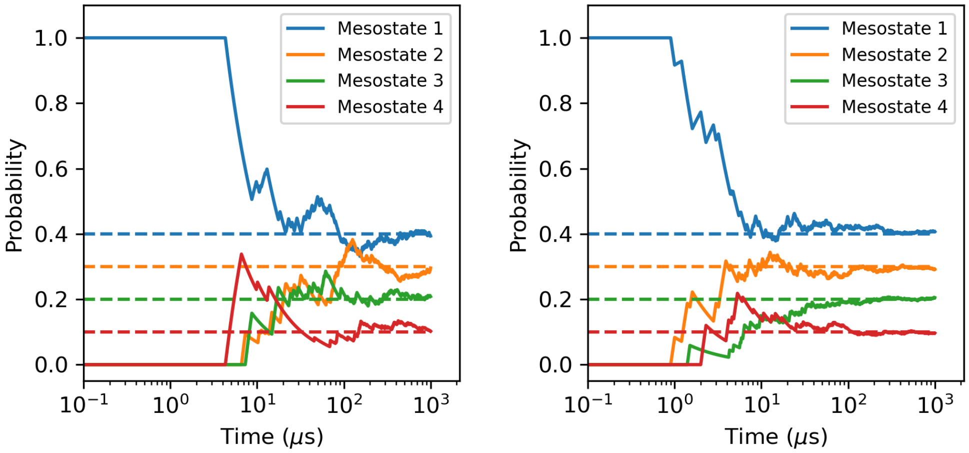


**Fig S2.** The evolution of mesostate occupation probabilities of a mechano-kinetic simulation of a simple dumbbell. Convergence to the mesostate averages is achieved whilst having explicit dependence upon the instantaneous mechanical energies due to thermal fluctuations. a) The slower set of rates. b) The faster set of rates.

### Coupled Model Validation

To observe how the mechanical energies directly affect the instantaneous transition rates, consider the following. If a thermal fluctuation causes a system within a particular mesostate to move closer to the mechanical equilibrium location in phase space of a different mesostate, then, as our mesostate transition protocol corresponds to no movement through phase space occurring with a kinetic transition, the mechanical energy change corresponding to a transition between those two mesostates will decrease. Hence, for a mechano-kinetic simulation that includes our energy-modified rates, we would expect the average mechanical energy change at the moment of kinetic transition (for a system with sufficiently large mechanical fluctuations) to be relatively low. On the other hand, if we included kinetic transitions without energy-modified rates, such that the instantaneous transition rates are at all times constant and equal to the average values defined in Table S2, we would instead expect the average mechanical energy change at the time of kinetic transition to be approximately equal to

$$\langle \Delta E_{ij} \rangle = \frac{1}{2} k_j \left( l_i - l_j \right)^2 , \tag{S1}$$

which is the mechanical energy change corresponding to a mesostate transition occurring at the equilibrium position of the initial mesostate $i$.

Fig.S3 shows the results of each type of simulation performed using the slower set of rates from Table S2, where Fig.S3a) has no energy-modified rates and Fig.S3b) includes our energy-modified rates as per Eqs.12 and 24. We see that in Fig.S3a), most of the transition energies are positive and relatively high compared relative to thermal energy. In a real physical system, kinetic transitions with no associated method of activation and such large increases in mechanical energy would be highly penalised. We see this penalisation in effect in Fig.S3b), where the explicit inclusion of the mechanical energies within each instantaneous transition rate causes transitions to occur only when that energy change is relatively small (or negative). While both types of simulation converge to the expected mesostate occupation probability distributions, it is clear that Fig.S3b) is more physically realistic for this system.

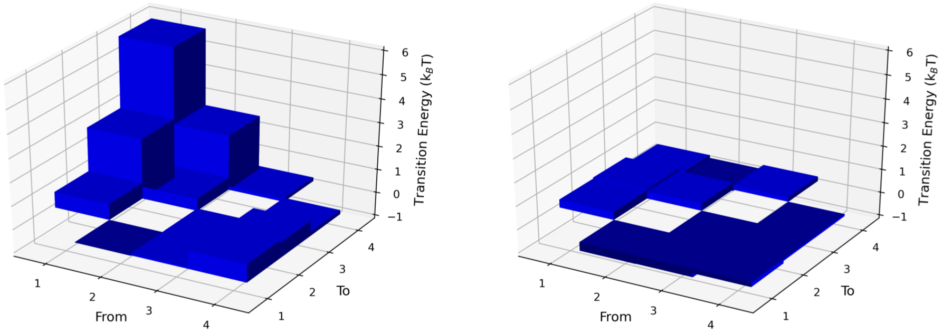


**Fig S3.** The effect of energy-modified kinetic transition rates on the coupling of dynamics and kinetics within the mechano-kinetic model. All leading diagonal values are zero as there is no mechanical energy change involved for a state remaining in the current state. a) No energy-modified rates. The lack of symmetry for forward and backward transitions is due to the differing values of $k_i$ in each mesostate (see Table S1) affecting Eq.S1. b) Energy-modified rates.
